# The Evolving Landscape of Neuroscience

**DOI:** 10.1101/2025.02.13.638094

**Authors:** Mario Senden

## Abstract

Understanding the evolution and structure of large scientific fields is crucial for optimizing knowledge production. Neuroscience is a rapidly expanding and diversifying field. To retain an overview of its cross-domain insights and research questions, this study leverages text-embedding and clustering techniques together with large language models for analyzing 461,316 articles published between 1999 and 2023 and reveals the field’s structural organization and dominant research domains. Inter-cluster citation analysis uncovers a surprisingly integrated picture and key intellectual hubs that shape the broader landscape. The field further exhibits a strong experimental focus, widespread reliance on specific mechanistic explanations rather than unifying theoretical frameworks, and a growing emphasis on applied research. Fundamental research is at the risk of decline and cross-scale integration remains limited. This study provides a framework for understanding neuroscience’s trajectory and identifies potential avenues for strengthening the field, offering a model for understanding the trajectory of complex research communities.

## Introduction

We have a longstanding desire to decipher the mystery of our own mind^1^. While philosophy and psychology have provided important insights regarding our cognitive and emotional makeup, neuroscience is unique in its ambition to provide mechanistic explanations of the mind^2^. Modern neuroscience, rooted in early work of Camillo Golgi and Santiago Ramón y Cajal^3,4^, has rapidly advanced through seminal work such as Sherrington’s description of reflexes^5^, Hebb’s theories on synaptic plasticity^6^, Hubel and Wiesel’s work on the visual cortex^7–9^, and the discovery of grid cells by Edvard and May-Britt Moser and their students^10–12^. Hand in hand with an ever-increasing pace of scientific discoveries also came diversification of the field into increasingly specialized research domains, such as work specifically devoted to the neural mechanisms underlying rare neurological syndromes (e.g.^13–16)^. While this is a natural tendency of scientific disciplines^17,18^, it can obfuscate the interconnectedness of phenomena and research questions and may thus hinder further progress^19,20^.

Consequently, there is an urgent need to provide a high-level perspective on the evolving landscape of neuroscience, both to integrate the field and to understand the dynamics driving a major area of human intellectual endeavor. Addressing this necessity, I have compiled an extensive dataset of neuroscientific abstracts and their metadata spanning from 1999 to 2023 from Q1 and Q2 journals in the respective years according to the SCImago Country & Journal Rank (SJR) database22. Utilizing state-of-the-art text embedding and clustering techniques, I categorized the literature into 175 distinct research clusters, enabling a content-driven examination of the field’s dynamic trajectory over the past 25 years.

Through these clusters, I systematically map the structural organization of contemporary neuroscience, identifying dominant themes as well as trends and open questions. By analyzing inter-cluster citation patterns, I reveal the extent to which different research domains interact, distinguishing between insular clusters that primarily cite their own work and integrative clusters that serve as intellectual bridges between domains. Furthermore, by characterizing clusters along key descriptive dimensions, I provide a detailed account of how neuroscientific research is organized. Detailed insights into individual research clusters, including their recent trajectories and major open questions, are presented in the Supplementary Materials. Together, this work offers a data-driven framework for understanding the evolution of neuroscience as well as a new dataset that can inform future research directions and promote new collaborative initiatives to bridge specialized domains and advance our collective understanding of the brain.

## Results

### Publication and Citation Patterns in Neuroscience

My first goal was to provide a snapshot of neuroscience publishing trends in the period from 1999 to 2023. To that end, I first identified all neuroscience journals ranked in the top two quartiles in the respective years according to SCIMago Journal Rank. I supplemented these neuroscientific journals with Q1 multidisciplinary journals that publish neuroscientific research such as Nature, Science and Plos ONE. The next step involved querying PubMed for a maximum of 5,000 articles per year for each selected journal. Through this procedure I obtained metadata and abstracts for research and review articles, excluding editorials and similar content. However, the procedure also yielded articles from disciplines other than neuroscience. To remove these, I embedded abstracts in a vector space using a general-purpose text embedding model from Voyage AI and created a custom discipline classifier neural network to filter out all non-neuroscience articles (see Methods for details).

This procedure resulted in a dataset of 461,316 neuroscientific articles published in 375 journals between January 1999 and December 2023^23^. The vast majority (88.16 %) of these are research articles. The total number of both types of articles that could be obtained from PubMed increased over time (Figure 1b), with a compound annual growth rate of 2.39% and 5.91% for research and review articles, respectively. While research articles heavily outnumber reviews, reviews exhibit higher citation rates, defined as the total number of citations divided by the article’s age. On average, research articles receive 4.5 citations per year (median = 2.5), whereas review articles receive 10 citations per year (median = 5.3; see Figure 1c for distributions of log-transformed citation rates). Interestingly, the median citation rates (MCRs) of both types of articles increased from 1999 to a peak in 2019, followed by a sharp decline. This rise and fall pattern in MCR indicates that many articles experienced a period during which they were most frequently cited sandwiched between an early phase when they were not yet receiving many citations and a later phase when they were no longer receiving many citations. Indeed, most citations to articles occur within one to 10 years following their publication with the peak typically at around three years (see Figure 1d,e).

**Figure 1:**
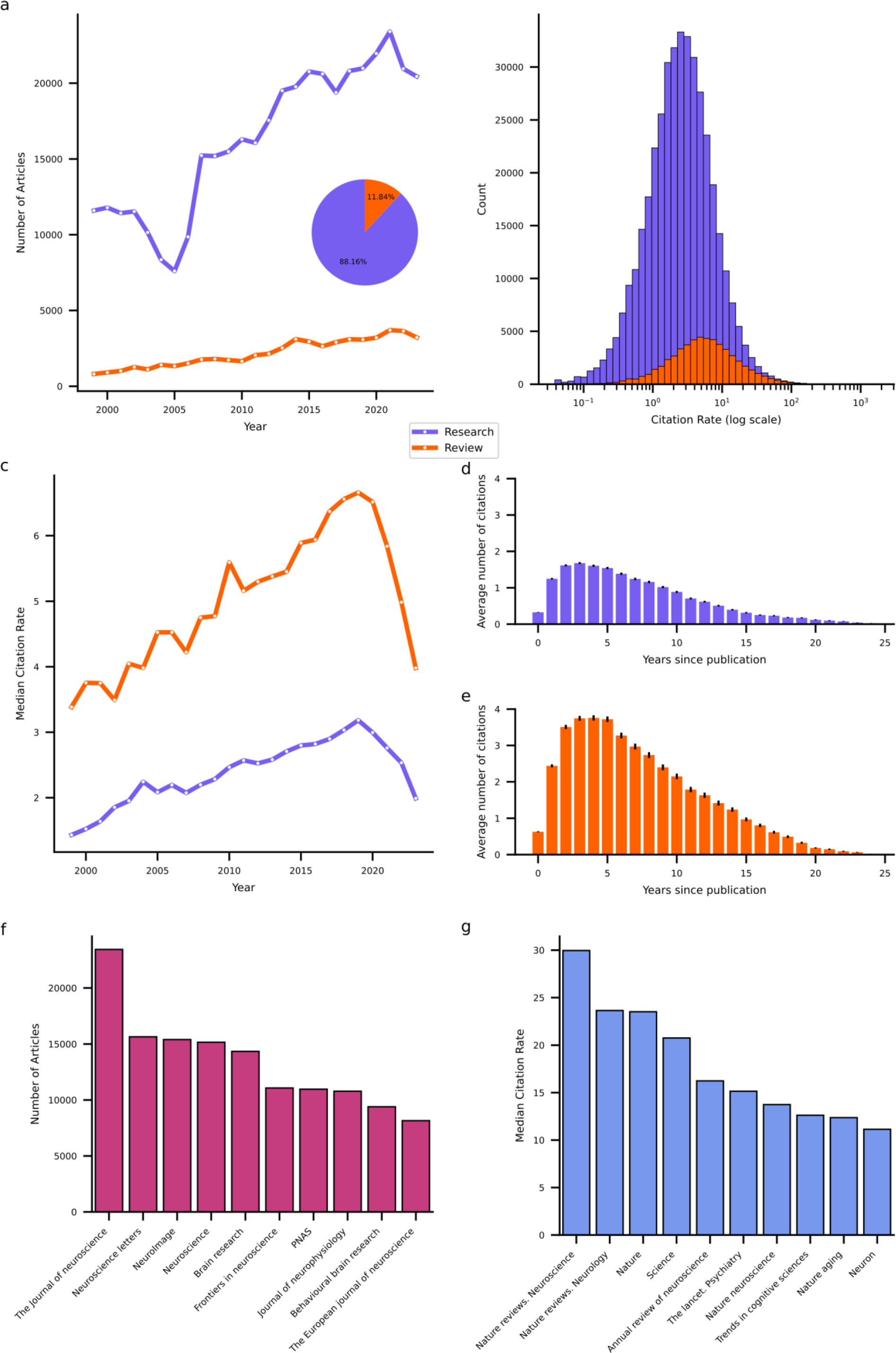
Publication and Citation Trends. **a**, Division between (purple) and review (orange) articles in the dataset broken down by publication year. Pie chart inset shows distribution for all years in aggregate. **b**, Distribution of log-transformed citation rates of research and review articles. **c**, Median citation rates of research and review articles by the year they were published. **d**, Average number of citations articles received as a function of their age for research articles. Estimated from a sample of 10,000 randomly chosen articles of each type published between 1999 and 2019. Error bars reflect standard error of measurement. **e**, Average number of citations articles received as a function of their age for review articles. Estimation procedure and error bars as in panel d. **f**, Number of articles in the dataset by the journal they were published in. Only the top 10 outlets are shown. **g**, Median citation rates of articles published in a given journal. Only the journals with the top 10 MCR values are shown.

To further characterize the publishing landscape, I identified the top 10 primary outlets that published the highest volume of neuroscience articles over the past 25 years (see Figure 1f) and the top 10 high-impact journals (see Figure 1g), defined here by the highest MCRs for articles published therein. The primary outlets list encompasses largely neuroscience specific journals that cater to a broad range of subdisciplines such as *The Journal of Neuroscience.* The list of high-impact journals likewise comprises domain specific journals with a wide scope. However, in contrast to the list of primary outlets, it contains a larger proportion of review-focused and high-profile multidisciplinary journals such as *Nature* and *Science*.

### Neuroscientific Research Domain Clusters

My second goal was to identify the distinct research domains that divide neuroscientific work. To that end, I clustered abstracts based on their semantic similarity measured as cosine similarity between their domain specific text embeddings. To achieve this, I first further embedded the abstract embeddings obtained with the general-purpose Voyage AI embedding model into a lower-dimensional, domain-specific latent space. I then constructed a semantic graph wherein each abstract is connected to its 50 semantically nearest neighbors, with each link weighted by the cosine similarity between its vertices. I then applied the Leiden community detection algorithm^24^ on this graph to obtain clusters. For each cluster, I submitted abstracts of the 200 nearest neighbors to the cluster’s centroid to a gpt-4o large language model (LLM) from OpenAI to describe the cluster. For clusters with less than 200 articles, I submitted all available abstracts. See Methods for details on these procedures.

Clustering applied to embedded abstracts identified 175 unique clusters ranging in size from 9,155 (cluster 0) to 117 (cluster 174) articles (Figure 2a). For a detailed overview of all clusters see Supplementary Table 1. The largest cluster involves research on the mechanisms of neuropathic pain including spinal cord modulation, glial activation, and receptor-mediated processes. The smallest cluster is concerned with the effects of electromagnetic fields emitted by mobile devices on brain function.

**Figure 2:**
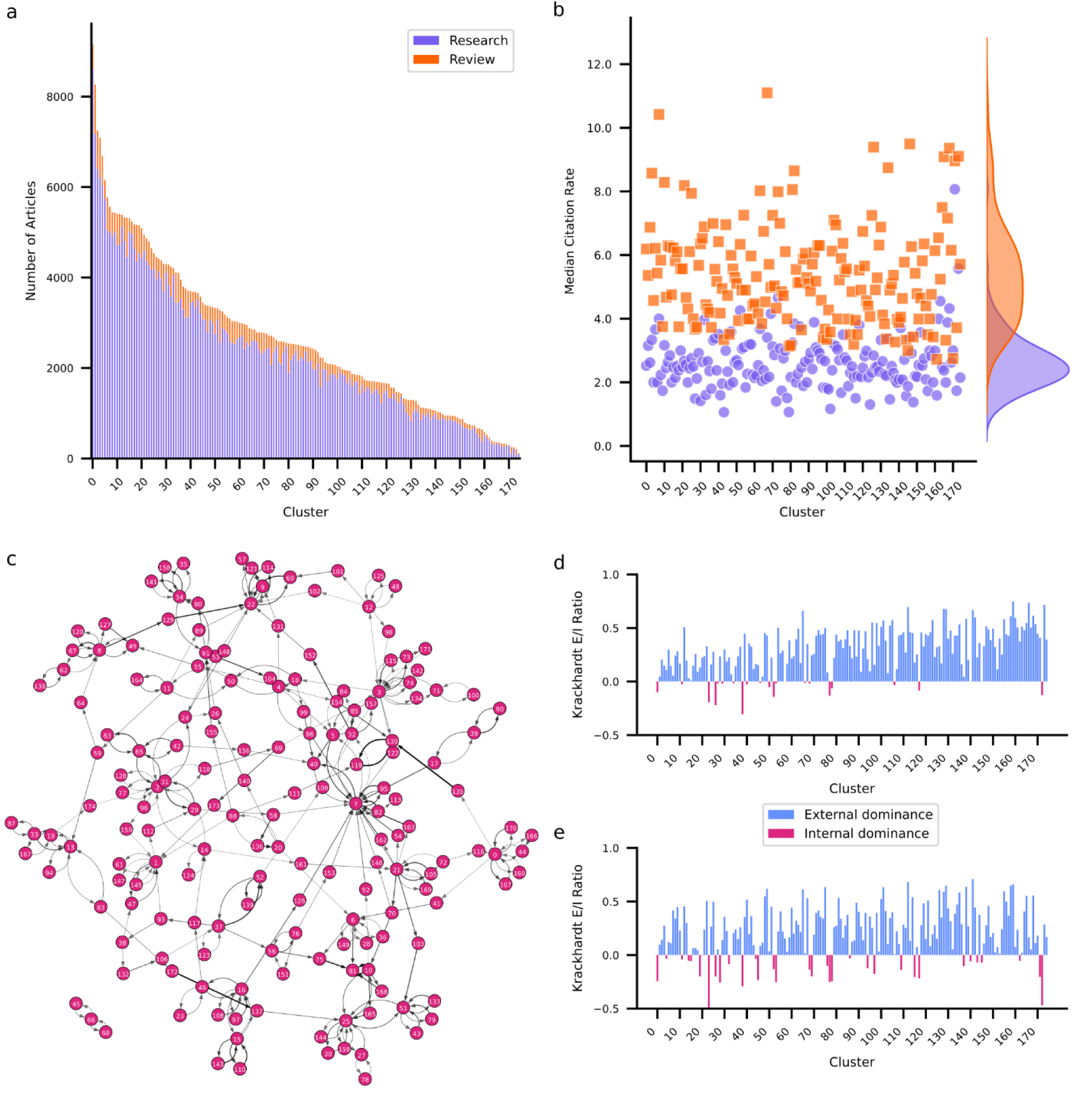
Research domain clusters. **a**, Number of articles within each cluster with review articles (orange) stacked on top of research articles (purple). **b**, Median citation rates of research and review articles per cluster. Distributions of MCRs are superimposed on the right. **c**, Cluster citation graph. Darker edges indicate higher citation density among articles within the connected clusters. **d**, Reference Krackhardt coefficient for each cluster. Positive values (blue) indicate that articles within a cluster primarily cite articles from other clusters. Negative values (red) indicate that articles within a cluster primarily cite articles in the same cluster. **e**, Citation Krackhardt coefficient for each cluster. Positive values (blue) indicate that articles within a cluster are primarily cited by articles from other clusters. Negative values (red) indicate that articles within a cluster are primarily cited by articles from the same cluster.

While most clusters are dominated by research articles, intriguingly two clusters contain more review articles. Cluster 171, which investigates the pathophysiology and potential long-term effects of SARS-CoV-2 on the nervous system, and cluster 173, which investigates the role of exosomes in neurodegenerative disease. Cluster 171 also presents the highest MCR for both research (8.1) and review (11.1) articles, likely explained by the immense interest in SARS-CoV-2 during the global pandemic^25–27^.

In terms of content, several clusters exhibit some degree of thematic overlap. A total of 14 clusters are devoted to Alzheimer’s disease (AD), each investigating distinct but complementary aspects. For example, cluster 1 investigates the role of amyloid beta peptides in the pathogenesis of AD whereas Cluster 47 focuses on the pathophysiological mechanisms involving tau protein modifications.

Furthermore, cluster 73 focuses on neuroinflammatory processes involved in AD and cluster 85 explores the use of cerebrospinal fluid biomarkers and neuroimaging techniques to advance diagnostic accuracy. This pattern is not unique to AD. There are nine clusters devoted to Parkinson’s, indicating that neurodegenerative diseases form their own group of clusters. Apart from clusters devoted to specific conditions, I also observed modality (e.g., vision and audition), cognitive/behavioral (e.g., decision-making, language, and memory), and methodological (functional neuroimaging, brain stimulation) groups of clusters. Curiously, my examination did not reveal any theory-specific clusters, suggesting that theoretical frameworks may be embedded within, rather than defining, research domains.

### Inter-Cluster Citation Structure

To further examine how clusters relate to each other, I examined the interactions between clusters in terms of their citation structure. To that end, I identified for each cluster which other cluster most frequently cites its articles, and which other cluster most frequently gets cited by its articles. I used this information to construct a citation graph among clusters (Figure 2c). Each edge in the graph is weighted by the citation density (fraction of articles that are cited to the number of articles that could be cited; see Methods for details). Additionally, I computed the Krackhardt coefficient^28^ for each cluster’s incoming links (citations) and outgoing links (references). For an overview of graph metrics per cluster see Supplementary Table 2. Notably, 74.86% of clusters exhibit positive Krackhardt coefficients for both citations and references. These clusters predominantly cite and are cited externally, suggesting diffusion of knowledge across clusters. These clusters lean on research from other clusters (indicated by a positive Krackhard coefficient for their references) but also provide insights for other clusters (indicated by a positive Krackhardt coefficient for their citations). Cluster 159, which focuses on the neural underpinnings of consciousness, is a notable example of an externally focused cluster. Conversely, only 6.86% of clusters contain articles that frequently cite and are cited internally, indicated by negative Krackhardt coefficients for both their reference and citation patterns. For example, Cluster 23, which studies the mechanisms and outcomes of auditory damage and repair, exhibits this pattern. Clusters such as these exhibit high internal cohesion and specialization but are siloed off from the remainder of the field. Some clusters exhibit mixed patterns. One such group (13.14%) maintains positive Krackhard coefficient for their references but negative Krackhardt coefficient for their citations. This suggests that while these clusters predominantly cite externally, their own articles are primarily cited internally. This pattern arises for clusters that rely on insights and methodologies from other clusters but are thematically niche. One example of this is cluster 28 which investigates the effects of adolescent binge drinking on brain development. The obverse are clusters that cite internally while being predominantly cited externally. I observed this pattern in 5.14% of clusters, but with negative Krackhard coefficients for references close to zero implying that these clusters have only a slight tendency towards citing internally. Cluster 40 is an example of this and focuses on advanced neuroimaging techniques for characterizing brain anatomy.

It is important to note that while the Krackhardt coefficient is useful for assessing whether a cluster is more inward- or outward-focused, it does not necessarily indicate the overall influence of a cluster. A cluster that is cited more frequently by articles outside its own domain may still not be a major hub if it lacks high overall citation counts. To gain a clearer picture of which clusters play a centralizing role, I examined key citation graph metrics, including in-degree (number of citing clusters), PageRank (influence within influential clusters), and betweenness centrality (how well a cluster bridges different research areas). Several clusters emerge as hubs within the neuroscience research network. Cluster 7 stands out as the most influential, with the highest weighted in-degree (0.0125), highest PageRank (0.079), and high betweenness centrality (10,789). Cluster 22 is another major hub, with a high in-degree (0.0057), PageRank (0.051), and betweenness centrality (14,899). Examining which clusters cite research in these clusters reveals that they are central because they provide complementary insights or relevant methodologies for other clusters. For example, cluster 22 (Molecular Mechanisms of Synaptic Plasticity in the Hippocampus) feeds into cluster 60 (Mechanisms and Dynamics of AMPA Receptor Trafficking and Synaptic Plasticity) and cluster 7 (Dynamic Functional Connectivity in Resting-State fMRI) feeds into cluster 5 (Functional Connectivity and Network Dynamics in Preclinical and Clinical Alzheimer’s Disease). By contrast, clusters 150 and 72 are highly isolated with low in-degree (0.00004 and 0.00009 respectively), PageRank scores (0.0010 and 0.0009) and betweenness centrality (2,158 and 2,212). These clusters exhibit highly specialized research focuses on thyroid hormones and vitamin D in neurodevelopment (cluster 150) and neuroendocrine and autonomic mechanisms in cardiovascular regulation (cluster 72) which likely restrict their integration into the broader neuroscientific landscape.

### The Dimensions of Neuroscientific Research

I followed these cluster-level analyses with a systematic investigation of the underlying dimensions that characterize neuroscientific research. To that end, I defined 10 dimensions (Appliedness, Methodological Approach, Species, Spatial Scale, Temporal Scale, Modality, Cognitive Complexity, Theory Engagement, Theory Scope, and Interdisciplinarity) and submitted abstracts of the 250 nearest neighbors to a cluster’s centroid to the LLM to characterize each cluster along these dimensions. A detailed assessment of the dimensions in each cluster is provided in Supplementary Table 3. Note that the categories that characterize a dimension are not mutually exclusive. A quantitative overview of how many clusters qualify for categories is shown in Figure 3a and reveals a predominantly experimental focus (96% of clusters) that employs both hypothesis-driven (77%) and data-driven (67%) research.

**Figure 3:**
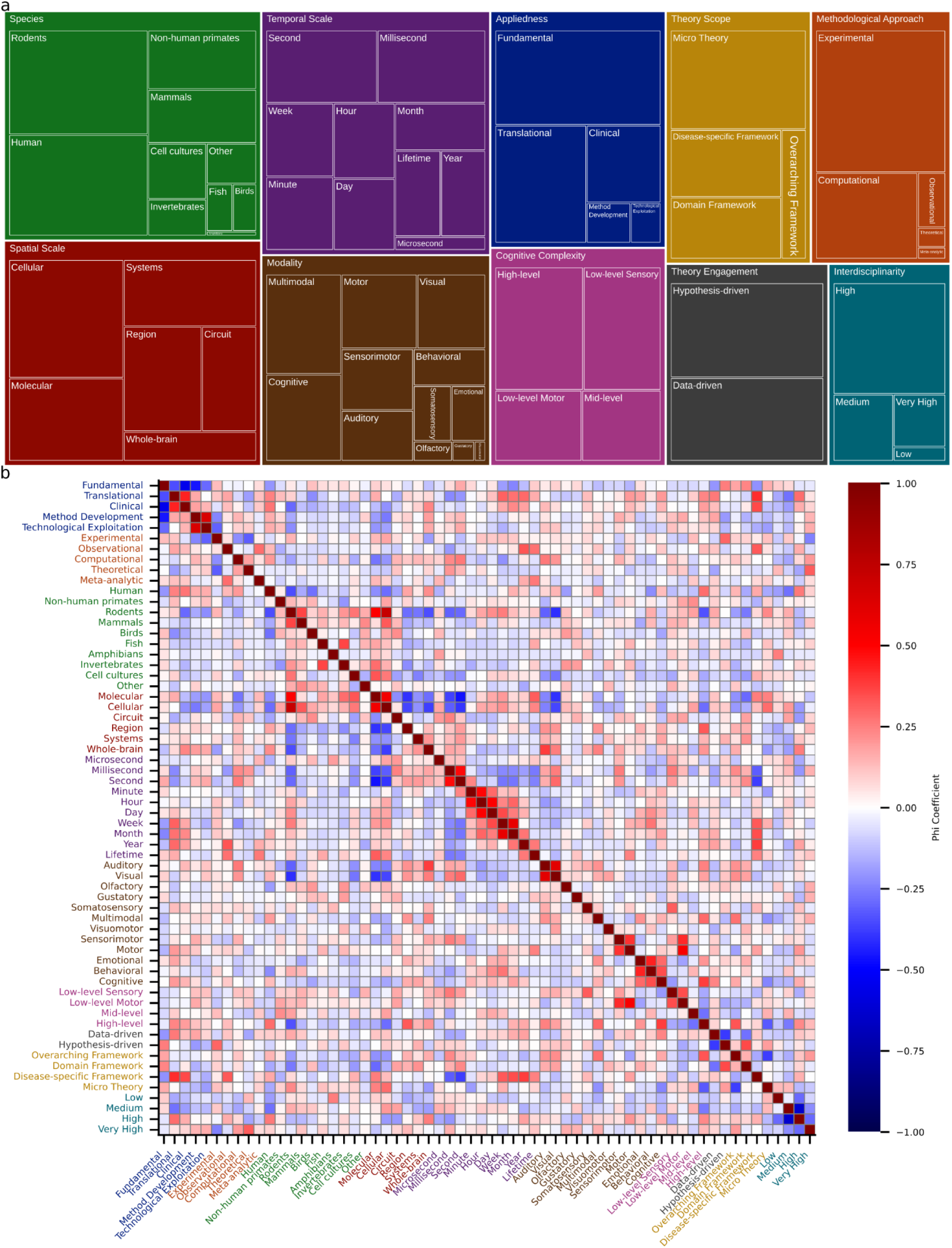
*Dimensions that characterize neuroscientific research. **a**, Tree map visualizing number of clusters qualifying for a given category within each dimension. Size of a region reflects the number of clusters that contain work exhibiting the category. Note that categories are not exclusive, and the same cluster may count towards several categories even within the same dimension. **b**, Phi correlation coefficients between all pairs of categories. Colors of category labels correspond to colors in panel a and indicate the dimension each category represents*.

Neuroscientific research often contains a theoretical element in the sense that it employs computational modeling (46%). However, only 3% of clusters contain work that develops, refines, or critiques conceptual frameworks and theories. This aligns with the observation that neuroscience tends to employ micro-theories (64%), i.e., narrowly scoped mechanistic accounts. Theories of intermediate scope in the form of domain-specific (35%) and disease-specific (39%) theories are also prevalent.

However, only 17% of clusters contain research that employs overarching theoretical frameworks aiming to explain fundamental principles of brain function. I observed that translational (55%) and clinical (30%) work sit atop a broad base of fundamental science (81%). Many clusters contain work on rodents (74%) or humans (71%), though non-human primates (33%) and other mammals (30%) also feature prominently. Neuroscience generally shows a balanced division between spatiotemporal scales, though work at the microsecond scale is sparse (6%). Finally, neuroscience displays high levels of interdisciplinarity, with 82% of clusters scoring high or very high.

I next sought to understand the extent to which categories co-occur by examining the Matthews correlation (phi) coefficient between pairs of categories (Figure 3b). The Matthews phi coefficient is the Pearson correlation coefficient estimated for two binary variables^29^. Please note that all reported p-values are Bonferroni corrected for multiple comparisons. Focusing first on spatial and temporal scales, it can be observed that molecular- and cellular-level investigations frequently go hand in hand (ϕ = 0.5502, t(173) = 8.667, p << 0.0001). By contrast, both molecular and cellular scales are negatively correlated with regional scale (ϕ = –0.4048, t(173) = –5.822, p << 0.0001 and ϕ = –0.3631, t(173) = – 5.126, p = 0.00148, respectively). The cellular scale also exhibits a negative correlation with the whole-brain scale (ϕ = –0.3773, t(173) = –5.359, p = 0.000497). Given that other correlations among spatial scales are not significant, this suggests a separation of the micro from the meso and macro scales. For temporal scales the interesting pattern emerges that neighboring scales are significantly correlated. For example, minute and hour exhibit a correlation of 0.4880 (t(173) = 7.353, p << 0.0001) and hour and day a correlation of 0.5142 (t(173) = 7.887, p << 0.0001). Only the consecutive pair of second and minute does not exhibit a significant correlation. This suggests that temporal scales are bridged through pairwise interactions rather than collectively. Interestingly, the molecular spatial scale is positively associated with the lifetime temporal scale (ϕ = 0.3144, t(173) = 4.356, p = 0.04262). Otherwise, fast temporal scales are negatively associated with small spatial scales (molecular – second: ϕ = –0.4640, t(173) = –6.890, p << 0.0001; cellular – millisecond: ϕ = –0.3131, t(173) = –4.337, p = 0.04622; cellular – second: ϕ = –0.3968, t(173) = –5.686, p = 0.000102; molecular – millisecond: ϕ = –0.3734, t(173) = – 5.294, p = 0.0006745).

Lastly, I examined how appliedness and theoretical scope interrelate. Notably, there is a sharp division between fundamental and all forms of applied research. Specifically, fundamental research is significantly anticorrelated with clinical (ϕ = –0.5496, t(173) = –8.6535, p << 0.0001), translational (ϕ = – 0.3441, t(173) = –4.8206, p = 0.00587), and method development (ϕ = –0.4553, t(173) = –6.7260, p << 0.0001) approaches. At the same time, translational and clinical research (ϕ = 0.4321, t(173) = 6.3024, p << 0.0001) and method development and technological exploitation (ϕ = 0.6530, t(173) = 11.3418, p << 0.0001) are positively associated. For theoretical scope, there is a negative correlation between micro theories and overarching theoretical frameworks (ϕ = –0.3752, t(173) = –5.3234, p = 0.000588). Perhaps unsurprisingly, both clinical and translational research is positively associated with work utilizing disease-specific theories (clinical: ϕ = 0.3539, t(173) = 4.9763, p = 0.00293; translational: ϕ = 0.4082, t(173) = 5.8821, p << 0.0001).

### Emerging Trends and Developments

My last goal was to identify trends in neuroscience. At the individual cluster level, I examined growth trends in terms of the size-adjusted annual growth rate, which quantifies the yearly increase in article count for each cluster relative to its total number of articles. Figure 4 summarizes the results and shows that a majority (52.0%) of clusters have increased their output above what is expected based on increases in output of the entire discipline. One third of clusters exhibit stable output as they neither decline nor exceed the growth of the discipline. It must be acknowledged that depending on their exact growth rate, these clusters may be considered to exhibit a relative decline as their growth is outpaced by that of the discipline. Finally, 15.4% of clusters exhibit an absolute decline in their output. Notably, the SARS-CoV-2 cluster (171) is among the ten fastest growing clusters. Generally, it appears that growing clusters share a strong applied focus and target overarching themes such as neurodegeneration (clusters 69 and 141), neuromodulation (clusters 167 and 168), and technological advancements in neuroscience (clusters 94 and 120). By contrast, declining clusters reflect predominantly fundamental research with a focus on receptor dynamics (clusters 9, 55, 60, 75, 89, 101, 124, and 135) and signaling pathways (clusters 48 and 102). These extremes of the spectrum reflect larger trends across clusters.

**Figure 4:**
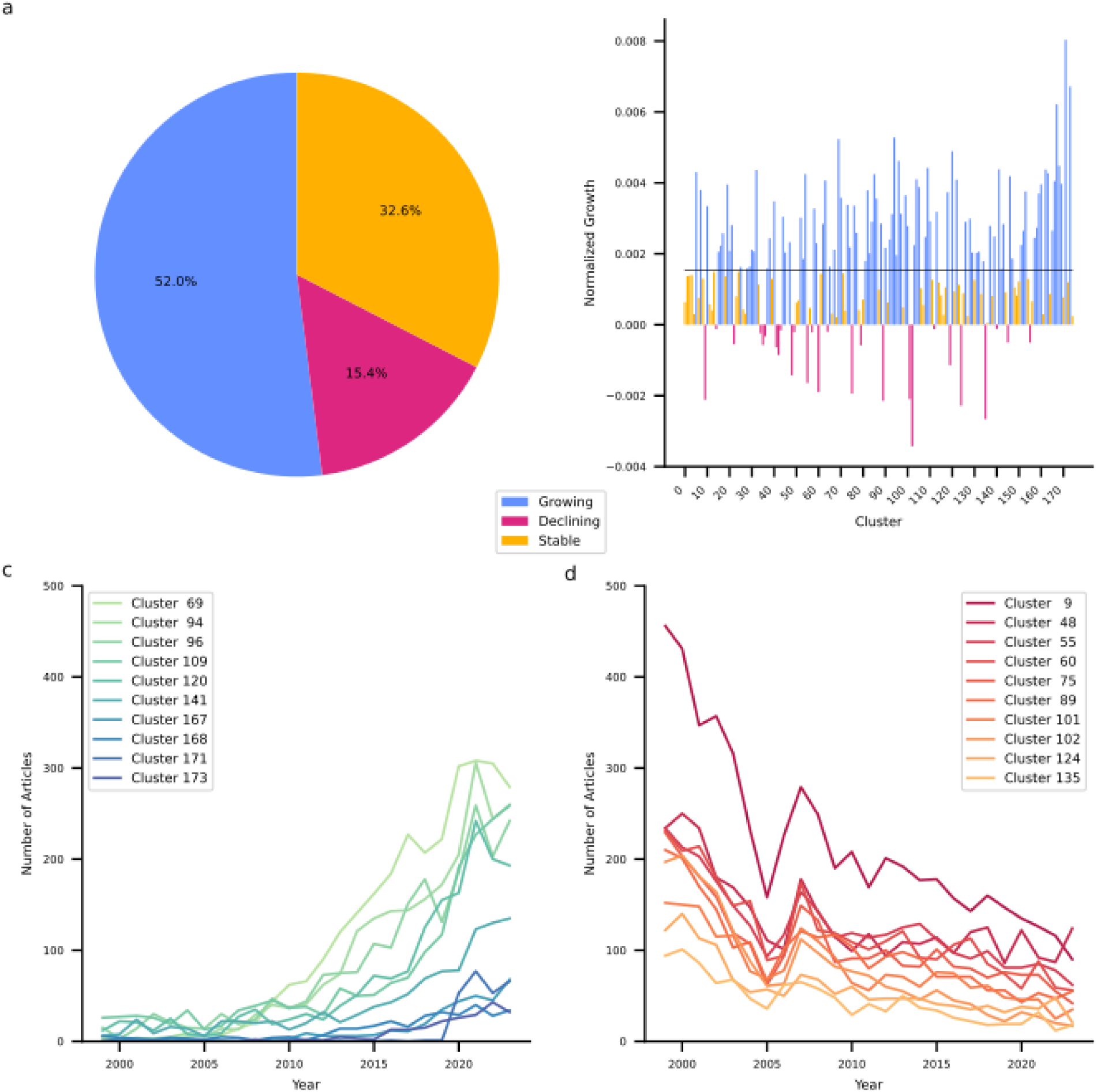
Quantitative growth trends. **a**, Fractions of clusters that exhibit growth (blue), decline (red), or remain stable relative to neuroscience as a whole. **b**, Normalized growth rates of each cluster. Colors as in panel a. The horizontal black line indicates the normalized growth rate of neuroscience. **c**, Number of articles in the dataset for the ten most growing clusters broken down by year. **d**, Number of articles in the dataset for the ten most declining clusters broken down by year.

While clusters containing fundamental research together are growing at a compound annual growth rate of 1.89%, this is less than the growth exhibited by neuroscience globally (2.39%). By contrast, clusters involving translational research, clinical research, method development, and technological exploitation exhibit compound annual growth rates of 3.57%, 4.78%, 7.51%, and 7.80%, respectively.

I followed up on these quantitative analyses with a more in-depth, qualitative, analysis of recent trends in neuroscience. First, I investigated the evolving themes within each research cluster by randomly selecting up to 100 articles published between 2010 and 2021, along with the top 100 cited articles from the same clusters since 2021. These selections were analyzed using an LLM with the objective of identifying both emerging and declining thematic and methodological trends. Furthermore, for each cluster, I curated five review articles published in 2023 (part of the dataset) or 2024 and concurrently submitted all full articles to the LLM to synthesize the major open questions highlighted therein.

Researchers in each domain can find specific current trends and key open questions in Supplementary Table 4. To develop a more holistic understanding of overarching trends, I aggregated all cluster-specific trends and questions and tasked the LLM with identifying overarching trends, transcendental questions, and necessary developments to move the field forward. These results are summarized in table 1.

**Table 1:** Emerging Trends, Key Scientific Questions, and Necessary Developments in Neuroscience.

| Overarching Trends | Transcendental Questions | Necessary Developments |
| --- | --- | --- |
| Integration of multi-omics and advanced imaging techniques in neuroscience research is gaining momentum, providing a comprehensive understanding of neurological disorders and cognitive functions by combining genomic, transcriptomic, proteomic, and imaging data. | How can novel methodologies like single-cell RNA sequencing and cryo-electron microscopy be used to uncover the precise mechanisms of synaptic plasticity and neuronal connectivity in various neurological disorders? | Integration of multi-modal imaging and sequencing technologies to understand complex brain network dynamics in real-world environments, highlighting the need to combine advanced neuroimaging with gene expression data. |
| There is an increasing focus on sex differences and individualized approaches in the study of neurological disorders and therapeutics, emphasizing the importance of personalized medicine and targeted interventions. | How do genetic and environmental factors interact to influence susceptibility to neurological diseases, and how can this knowledge inform personalized medicine approaches? | Development of comprehensive, personalized therapeutic interventions targeting neurodegenerative and psychological disorders by leveraging insights from genetic, environmental, and neuroinflammatory factors. |
| The use of machine learning and artificial intelligence in analyzing complex datasets such as neuroimaging, electrophysiological recordings, and behavioral data is enhancing the precision of diagnostic and therapeutic strategies across various neurological and psychiatric conditions. | In what ways can artificial intelligence and machine learning be integrated with neuroimaging and 'omics' data to enhance our understanding of brain disorders and improve diagnostic and therapeutic approaches? | Integration of AI and machine learning across neuroscience research to improve diagnostic accuracy and develop predictive models for neurological and psychiatric condition progression, incorporating multi-modal datasets. |
| Neuroinflammation and immune pathways are recognized as critical components in the pathogenesis and progression of neurodegenerative diseases, highlighting novel therapeutic targets and the importance of glial cell functions. | What roles do neuroimmune interactions and inflammation play across different neurodegenerative and psychiatric disorders, and can they be harnessed for therapeutic interventions? | Advancement in the understanding and therapeutic targeting of neuroimmune and neuroinflammatory pathways in diseases like Alzheimer's, which requires integrating genomic data with inflammation and gut-brain axis research. |
| The importance of neuroplasticity and the dynamic reconfiguration of neural networks is emphasized in understanding cognitive processes, learning, memory, and recovery from brain injuries, offering new insights into adaptive neural changes. | What are the mechanisms behind neuroplasticity and connectivity changes in response to both pathological and therapeutic interventions, and how can this knowledge be applied to enhance cognitive and motor recovery? | Expansion of high-resolution and non-invasive brain imaging technologies to better capture and predict neurodegenerative and neuropsychiatric disease progression in personalized medicine approaches. |
| Emerging non-invasive and precise neuromodulation techniques, such as ultrasound and optogenetics, show promise in modulating neural circuits for therapeutic interventions in neurological disorders, focusing on targeted and adaptable treatment strategies. |  | Breakthrough in targeted modulation of specific brain circuits using non-invasive technologies like focused ultrasound, emphasizing the need for precision in neuromodulation therapies for conditions like depression and epilepsy. |
| Neurodevelopmental and neurodegenerative diseases are increasingly being studied through advanced models like organoids, induced pluripotent stem cells, and detailed human-specific studies, refining insights into disease mechanisms and therapeutic approaches. | How can single-cell and multi-modal data from transcriptomics, proteomics, and imaging be combined to elucidate cell-type-specific roles in neural development and neurodegeneration? | Innovation in regenerative medicine and organoid technology to provide new insights into human developmental disorders and neurodegeneration, using advanced materials and engineering methods. |
| Research into the gut-brain axis and its influence on neurodegenerative and psychiatric disorders is expanding, | How do gut-brain interactions contribute to the onset and progression of neurodegenerative | Comprehensive modeling of the gut-brain axis to uncover its involvement in neurological health and disease |
| exploring the role of microbiota and gut-derived metabolites in brain health and disease, with potential for microbiome-based interventions. | diseases, and what therapeutic interventions can modulate these pathways for disease prevention and treatment? | states, particularly in the context of neurodegenerative and psychiatric disorders. |
|  | What are the signaling pathways involved in the neuroprotective mechanisms against stress and trauma, and can they be modulated for therapeutic benefits in neuropsychiatric conditions? |  |
| Astrocytes and glial cells are increasingly recognized for their active roles in neurophysiology and neuroprotection, influencing synaptic plasticity, neuroinflammation, and disease progression, representing potential therapeutic targets for CNS disorders. | How can advances in real-time and multi-modal imaging technologies deepen our understanding of dynamic brain processes and guide the development of effective interventions for cognitive and behavioral disorders? | Elucidation of the mechanistic role and therapeutic potential of astrocytes and microglia in synaptic plasticity and neurodegenerative conditions, using advanced tools like optogenetics and CRISPR for functional insights. |
| Neurosymbolic integration and neuromorphic computing are advancing in neuroscience, emulating biological brain functions in artificial systems and enhancing understanding in cognition and decision-making processes. |  |  |
|  | What role do epigenetic modifications and regulatory RNA molecules play in the pathogenesis of psychiatric and neurodegenerative disorders, and how can these be targeted for novel therapies? |  |
|  |  | Improvement in the understanding of synaptic interaction mechanisms using advanced imaging and computational tools, focusing on neurotransmitter dynamics and synaptic plasticity in health and disease. |

Trends, questions and developments are predominantly aligned with applied research. This is consistent with the growth patterns of the research clusters, where applied areas are experiencing increased output. Notably, there is an absence of calls to develop or test theoretical frameworks. This does not imply that such efforts are absent in neuroscience, but they are less prominent relative to other trends.

## Discussion

The present study provides a comprehensive, data-driven mapping of the evolving landscape of neuroscience, serving as a large-scale analysis of knowledge production and community structure within a major scientific field. Neuroscience publishing has expanded steadily^21^ with broad neuroscientific journals as primary outlet. Research articles consistently outnumber reviews. The latter, however, exhibits higher citation rates. The journals with the highest impact thus unsurprisingly comprise review-focused neuroscientific journals, but also high-prestige multidisciplinary journals. A clustering analysis identified 175 distinct research domains with diverse themes such as neuropathic pain, neural underpinnings of consciousness, and EEG-based brain-computer interfaces. While several clusters are devoted to specific diseases, modalities, methods, and cognitive function, notably not a single cluster is dedicated to a theoretical framework.

An analysis of the cluster-level citation network revealed that most clusters integrate and spread insights from diverse research domains. Key hub clusters play a central role in shaping neuroscience by providing methodological and conceptual foundations for other clusters. Some clusters, however, are highly specialized and remain largely isolated from the broader research landscape. The dimensional analysis of neuroscientific research reveals that the field is predominantly experimental, with both hypothesis-driven and data-driven approaches. While computational work is prevalent, theoretical work remains limited. Furthermore, the field tends to employ micro theories more than overarching theoretical frameworks. This observation aligns with the lack of theory-focused clusters. Additionally, neuroscience exhibits clear structural divides, including a sharp distinction between fundamental and applied research and a separation of micro- and macro-level spatial scales. Interestingly, temporal scales are pairwise, but never fully integrated. Finally, a trends analysis revealed that neurosciences exhibit a shift towards more applied and translational research, with growing clusters focusing on neurodegeneration, neuromodulation, and technological advancements, while traditionally dominant fundamental research areas, such as receptor dynamics and intracellular signaling pathways, are in decline. The rapid growth of the SARS-CoV-2 cluster indicates that neuroscience is quick to respond to global events.

Overall, neuroscience appears to be thriving as it maintains high output across diverse topics ranging from neurodegenerative diseases, neuromodulation, cognitive functions, to technological advances. Despite this diversity, neuroscience achieves a high level of integration and extensive knowledge exchange across its domains. Targeted collaborations across clusters could further enhance this integration. The field currently exhibits a good balance between hypothesis-driven and data-driven approaches and between fundamental and applied research. However, growth trends show that fundamental research is losing ground. The underlying reasons for this remain unclear. It could reflect a change in funding priorities, where agencies increasingly emphasize clinical and technological applications. Alternatively, it may indicate a natural progression, where certain foundational questions have been sufficiently addressed, allowing researchers to pivot more towards applications. Regardless of the cause, this trend raises important questions about the long-term trajectory of neuroscience. If the shift toward applied research continues, will neuroscience still generate foundational discoveries at a sufficient rate? To address this, future studies could systematically assess the distribution of funding across fundamental and applied domains. In the meantime, funding agencies should establish dedicated initiatives supporting fundamental research to prevent erosion of the foundational knowledge upon which future breakthroughs depend.

Neuroscience spans all levels of organization, from molecular and cellular studies to whole-brain dynamics. However, integration across spatiotemporal scales remains limited. There is, for instance, a clear divide between small and large spatial scales. Additionally, temporal scales are only integrated in a pairwise manner, leaving connections between different timescales indirect. An important question is whether these divisions reflect a meaningful organizational principle of the brain. Processes unfolding at certain scales may simply not be relevant for understanding processes at other scales as each constitutes a self-contained system whose governing principles, despite emerging from smaller scales, nevertheless operate independently^30,31^. For example, understanding network dynamics in large-scale brain activity may not require understanding how they arise from individual neuronal processes. If this is the case, neuroscience may need to develop better theoretical models to formalize and justify these separations. Alternatively, the divide between research at different scales may simply be a byproduct of disciplinary specialization. In that case neuroscientists should explore systematic ways to more comprehensively bridge spatial and temporal scales. The development of standardized methods to analyze multi-modal datasets at different levels of neural organization will be essential for this^32–34^. Additionally, computational modeling may offer a means to connect findings across scales and to reveal unifying principles^35,36^, or alternatively to help formalize and justify the separation of scales as an inherent principle of brain organization.

A particularly concerning revelation of this study is the field’s predominant reliance on highly specific micro theories rather than broader theoretical frameworks. While computational models are widely used to test mechanistic hypotheses, theoretical work that develops and refines overarching frameworks is notably scarce. Lacking such frameworks can impede collaboration and data integration across research domains and hence diminish the field’s ability to unify diverse phenomena and findings into a comprehensive understanding of the brain^37–39^. To address this neuroscience could benefit from explicit efforts to determine whether different micro theories represent truly distinct explanations or special cases of more general frameworks. The dataset compiled in this study together with the ability of LLMs to analyze vast amounts of text can provide invaluable assistance in this effort^23,40,41^. Such an AI-assisted approach may reduce the challenge of synthesizing highly diverse findings inherent in rapidly evolving fields. The dataset also enables researchers to trace the evolution of existing frameworks, such as predictive coding or the global neuronal workspace theory, to identify how they integrate into different research domains. At the same time, it needs to be acknowledged that no single overarching theory might be able to unify all levels of neural organization. Funding agencies should support theoretical initiatives that aim to answer this question or reflect explicit efforts to unify disparate mechanistic accounts, and theoretical neuroscience must be recognized as an essential component of the field.

In conclusion, neuroscience is a thriving and interdisciplinary field, but recognizing and addressing its structural limitations is crucial for future progress. The findings of this study highlight that ensuring that fundamental research remains valued, greater integration across spatiotemporal scales, and increased theoretical synthesis could strengthen the field. If the observed separations across spatial and temporal scales are not merely practical limitations but reflect an inherent organizational principle of the brain, then neuroscience may need to reconsider its theoretical foundation. Rather than seeking a single all-encompassing framework, a more fruitful approach may be to explicitly acknowledge that different levels of neural organization operate under distinct governing principles. In this perspective theoretical frameworks should clearly define the scale at which they apply, and research efforts should prioritize understanding the mechanisms that allow for scale separation, as well as instances where cross-scale interactions are essential. Beyond neuroscience-specific insights, this work contributes to the broader field of meta-science or the science of science. The methods employed demonstrate the power of computational techniques for mapping scientific terrains, while the observed patterns likely resonate with the evolution of other complex scientific disciplines. Understanding these general dynamics is critical for effective science policy and research management.

## Methods

All analyses were performed in a Python 3.12 environment managed by conda on a HP Z440 Workstation (Intel® Xeon® E5-1650 v4 CPU @ 3.60 GHz, 12 cores, 31 GB RAM), running Ubuntu 22.04.5 LTS. I used PyTorch (version 2.2.0) for building and training all neural networks on a GeForce GTX 1080 Ti graphics card using CUDA version 12.2.

### Data Source and Extraction

The PubMed database (www.ncbi.nlm.nih.gov/Pubmed) was used as data source for obtaining abstracts and metadata of neuroscientific articles published between 1999 and 2025. I used the biopython (version 1.81) package to programmatically query the database with the following search query: **(“{journal}”[Journal]) AND ((“{year}/01/01”[Date - Publication]: “{year}/12/31”[Date - Publication]))**. Here *journal* and *year* were variables that were assigned based on yearly journal ranks provided by the SCImago Country & Journal Rank (SJR) database^22^. Specifically, for each year from 1999 to 2023, I queried all journals ranked within the first two quartiles of the neuroscience rankings provided by SJR. Additionally, for each year, I queried all journals within the first quartile of multidisciplinary rankings provided by SJR. These were included because multidisciplinary journals such as Nature and Science are valuable outlets for neuroscientific research. Finally, I queried journals that ranked in the first quartile for all other disciplines defined by SJR for the years 2000, 2010, and 2020. These were included to train a discipline classifier (see below). For each query, a maximum of 5,000 PubMed entries was obtained.

Queries were performed between April 15^th^ 2024 and April 18^th^ 2024. For each article resulting from PubMed queries, I extracted its PubMed ID, its digital object identifier (doi), the journal it was published in as well as the month and year of publication, its title, abstract, its type. I rejected all articles that were not of research or review type and removed all articles whose abstracts had less than 50 or more than 500 words. Between December 27^th^ 2024 and January 3^rd^ 2025, I obtained citation counts for articles in the dataset, ensuring that each article had at least one full year for accumulating citations. Based on the publication date, I computed article age in years with respect to January 3^rd^ 2025 and used this age to compute citation ratios as the citation count divided by age.

### Data Filtering

The journals used in my queries do not exclusively publish neuroscientific work. To reduce the dataset to neuroscientific articles, I used a neural network trained on a multi-label discipline classification task. The discipline classifier network is a fully connected neural network. The architecture consisted of an input layer receiving 1024-dimensional text embeddings, two hidden layers with 256 and 64 exponential linear units (ELUs), and an output layer with 26 sigmoid-activated units, each corresponding to a discipline. To prevent overfitting, a dropout layer with a probability of 0.5 was applied before the final classification layer. The input embeddings were generated using the *voyage-lite-02-instruct* model from Voyage AI, a general-purpose text embedding model.

To train the discipline classifier, a labeled training dataset was constructed. First all articles from non-neuroscientific journals were included. Then 5% of all articles in journals publishing neuroscience were included. Labels for each article were obtained from the discipline assessment provided by SJR for each journal. Importantly, these labels are accurate at the journal level but not at the article level. A particular article published in a journal that publishes neuroscience but also other subdisciplines of life science, might nevertheless be a pure neuroscientific article. For this reason, training was performed in three stages. In a pre-training phase, the classifier was trained for 25 epochs on a subset of the training data limited to articles from journals that exclusively publish in a single discipline.

Following pretraining, the classifier was applied to a broader set of articles, including those from multidisciplinary journals. All discipline labels falling below 99% of the maximum class probability were removed. Note that this does not remove any articles; rather, only discipline labels were discarded while simultaneously ensuring that each article retained at least one label. The classifier was then trained for 50 additional epochs, incorporating these high-confidence self-labeled samples. In the final stage, the model was fine-tuned for 10 additional epochs using again only articles from journals that exclusively publish in a single discipline. In all stages, data was subdivided into training (80%), validation (10%), and test (10%) subsets. No explicit class balancing techniques, such as resampling or weighting in the loss function, were applied. Training in all stages was conducted using binary cross-entropy loss, an Adam optimizer with a learning rate of 0.0002, and a batch size of 64.

After training, the labeled articles used for training were assessed for inclusion in the final neuroscience dataset. The articles originally selected from exclusively neuroscience journals were automatically retained since they were already part of the intended dataset. For each article published in journals that do not exclusively publish neuroscientific work, I used the discipline classifier network to identify whether neuroscience was within 80% of the most probable discipline. This threshold was chosen to ensure that only articles with a strong neuroscientific focus were included while minimizing the risk of discarding relevant research. Articles that did not meet this criterion were removed.

### Semantic Clustering

To identify domain clusters in neuroscience, I initially embedded abstracts of all articles in the final dataset in a neuroscience-specific latent space to be able to assess semantic similarity between articles. This proceeded in two stages. First, I embedded all abstracts using the general-purpose *voyage-large-02-instruct* text embedding model from Voyage AI. Then I further embedded the resulting vector representations using a custom neural network to obtain neuroscience-specific representation of article abstracts. The network comprised three fully connected layers with progressively smaller dimensions, reducing the input 1024-dimensional embeddings to a 64-dimensional domain-specific embedding space. Batch normalization was applied after the first and second fully connected layers to improve training stability and accelerate convergence. An ELU activation function followed each of these layers to introduce non-linearity, and dropout was applied immediately after the ELU activation in both the first and second hidden layers, with a dropout rate of 0.1 to mitigate overfitting. The output of the final layer was L2-normalized, ensuring that all embeddings had unit length.

The network was trained using a contrastive learning approach with an N-pair loss function. The objective was to learn a representation where semantically similar articles were mapped closer together while dissimilar articles were pushed apart. To define training pairs dynamically, cosine similarity was computed between the general-purpose embeddings that served as input to the network. Pairs of articles with an initial embedding cosine similarity of at least 0.85 were designated as positive pairs, while pairs with a similarity below 0.75 were assigned as negative pairs. Intermediate cases, where similarity fell between these thresholds, were excluded from training. Similarity cutoffs for defining positive and negative pairs were defined based on domain knowledge and manual inspection.

Specifically, I computed the cosine similarity between the embedded representations of a set of abstracts that I had judged to be part of the same research domain and compared this to the similarity between abstracts that I classified as belonging to different subdomains. A substantial portion of the dataset, approximately 95%, served as training dataset. The remaining 5% of the dataset was reserved for validation to monitor convergence and prevent overfitting. The training dataset was used to optimize the embedding network with an Adam optimizer, an initial learning rate of 0.0001, and an L2 weight decay of 0.01 to improve generalization. A batch size of 256 was used. The learning rate was scheduled to decrease exponentially over the course of 2000 epochs, with a decay factor of 0.995 per epoch.

Additionally, an L2 regularization weight of 0.01 and a correlation weight of 0.1 were applied to further constrain the learned representations and promote meaningful structure in the latent space. To accelerate training convergence and improve generalization, I incorporated Grokfast^42^, a method that amplifies slow gradients during optimization, with a smoothing coefficient of 0.98 and an amplification factor 2.

Following training, the learned embeddings were used to construct a similarity graph using python-igraph (version 0.11.8) of neuroscientific research, where each node represents an article and edges between nodes are weighted by their cosine similarity. A K-NN search was performed using FAISS (faiss_cpu version 1.8.0.post1) to retrieve the top 50 most similar neighbors for each article and add these as edges to the graph. This results in a directed graph because an article might be among the top 50 nearest neighbors of another article, but not vice versa. I symmetrized the graph by adding bidirectional edges.

The resulting undirected, weighted graph served as the foundation for clustering neuroscience into distinct research domains using Leiden community detection^24^ implemented in the leidenalg package (version 0.10.2). Leiden clustering was performed using the Constant Potts Model (CPM)^43^, which allows for resolution tuning to optimize the balance between intra-cluster density and inter-cluster separation. To systematically determine the optimal resolution parameter, clustering was conducted across 1000 resolution values, evenly spaced between 0.001 and 1.0. To identify the optimal resolution, modularity scores were monitored, and clustering was halted early if modularity exhibited a consistent downward trend across five consecutive resolution steps. The best resolution parameter was selected as the value that maximized modularity across the examined range. Articles were then assigned to clusters based on the optimal Leiden partition, and these cluster assignments were mapped back to the original dataset.

### Citation Density Graph

To construct a citation density graph at the cluster level, I first extracted citation and reference relationships for each article in the dataset. For each article, I retrieved the PubMed IDs of all articles citing it (incoming citations) and all articles it cited (outgoing references). Only linked PubMed IDs present in the dataset were retained. I next computed citation density between all pairs of clusters by identifying all articles that cite across clusters and normalizing this by the number of possible citations, taking the relative article ages into account. Additionally, for each cluster I identified the clusters whose articles most frequently cite articles in the clusters and whose articles most frequently get cited by articles in the cluster. Finally, I constructed a weighted and directed citation density graph by connecting each cluster to their most citing and most cited clusters and weighing these connections by the citation density.

## Metrics

### Compound Annual Growth Rate

The com =pound annual growth rate over the period from *t*_0_ is given by 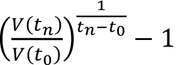, where *V*(*t*_0_) is the is the initial value and *V*(*t_n_*) is the final value.

### Size-Adjusted Annual Growth Rate

The size-adjusted annual growth rate is given by 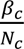, where *β_c_* is the slope obtained from a linear regression of the number of articles in cluster *c* as a function of years and *N_c_* is the total number of articles in cluster over the entire period.

### Krackhardt Coefficient

The Krackhardt coefficient (E/I-ratio) is given by 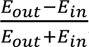, where *E_out_* is the number of external citations and *E_in_* the number of internal citations.

### LLM-Based Analyses

I conducted several semantic analyses of the domain clusters with the aid of OpenAI’s *gpt-4o-2024-08-06* large language model (LLM). First, I obtained a title, short description, relevant keywords, and a classification of the primary focus (thematic or methodological) for each cluster by submitting up to 200 abstracts of each cluster to the LLM. For clusters with less than 200 articles, I submitted all abstracts. For clusters with more than 200 articles, I selected those whose abstracts were most similar to the cluster centroid in the previously computed semantic embedding space. The selected abstracts were formatted and submitted to the LLM using the LangChain API (version 0.3.19 of langchain and 0.3.6 of langchain_openai), with the following prompt:

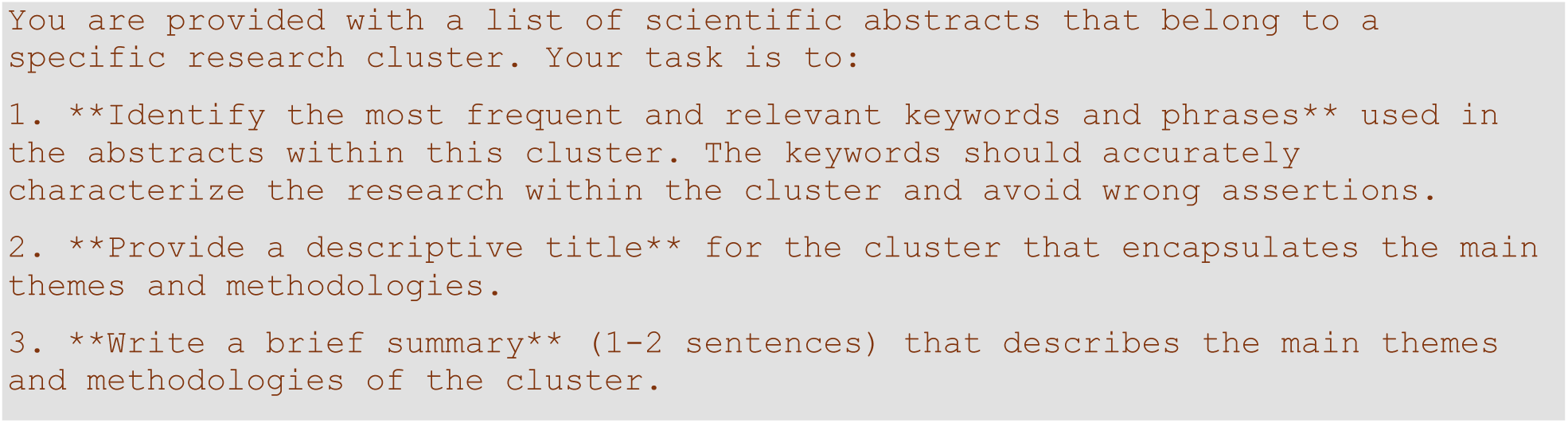
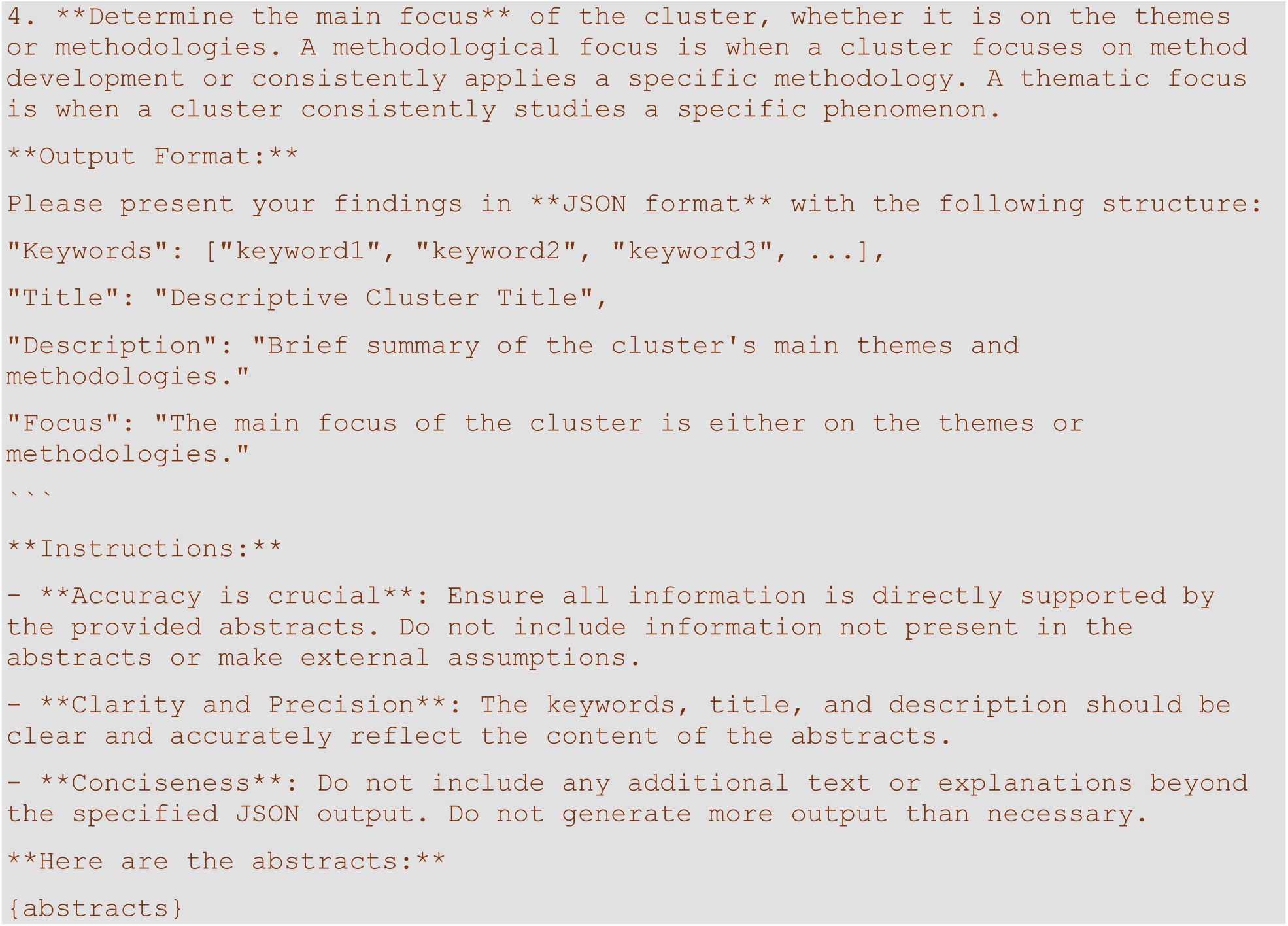

The generated JSON data was automatically extracted and integrated into the dataset. The prompt was designed to minimize the risk of hallucination or speculative assertions by the model, as it was explicitly instructed to rely solely on the provided abstracts.

To better understand what distinguishes similar clusters, I identified the most similar cluster for each of the 175 clusters. For each cluster, I then submitted the 100 articles that were most similar to the other cluster’s centroid as well as the 100 articles of the other cluster that were most similar to its centroid to the LLM with the following prompt:

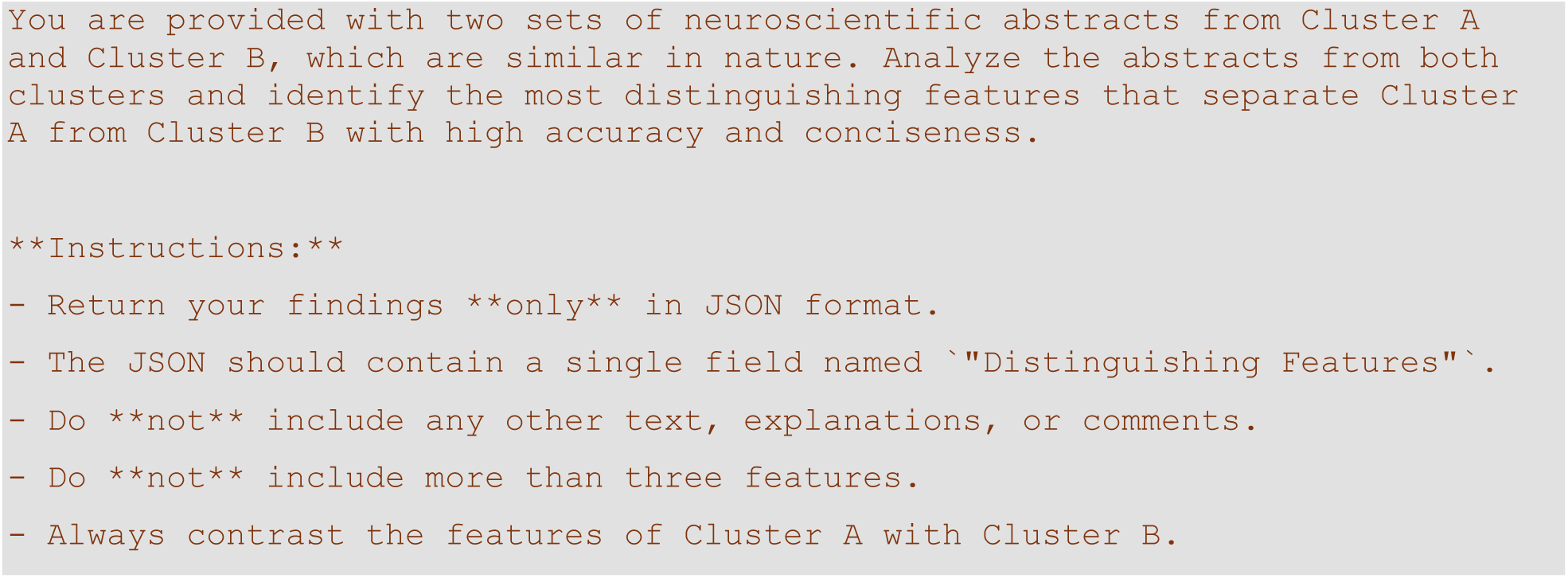
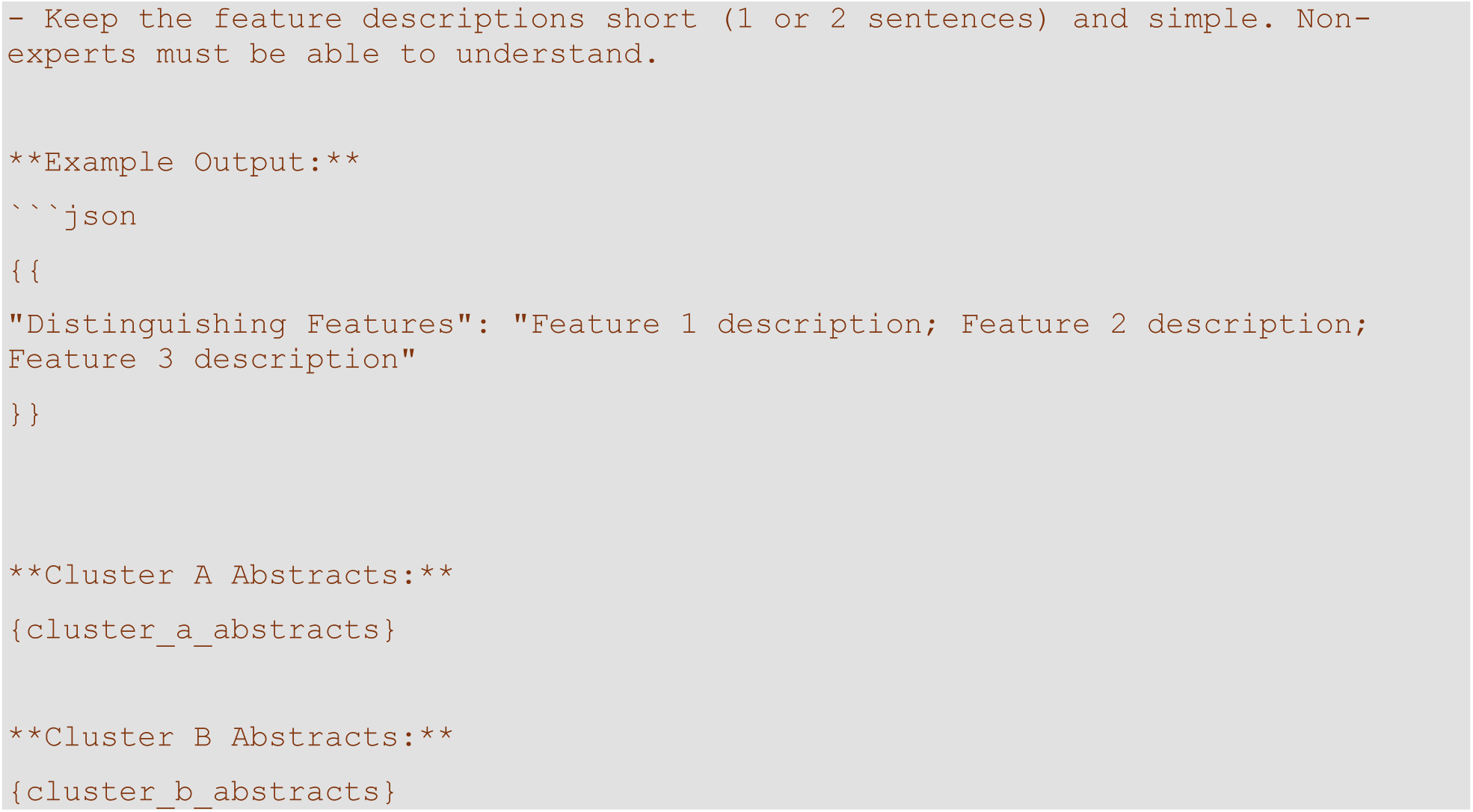

The generated JSON data was automatically extracted and integrated into the dataset.

Subsequently, I utilized the LLM to assess research within each cluster along 9 research dimensions: appliedness, modality, spatiotemporal scale, cognitive complexity, species, theory engagement, theory scope, methodological approach, and interdisciplinarity. To that end, I submitted up to 250 abstracts of each cluster together with the cluster title to the LLM using the following prompt:

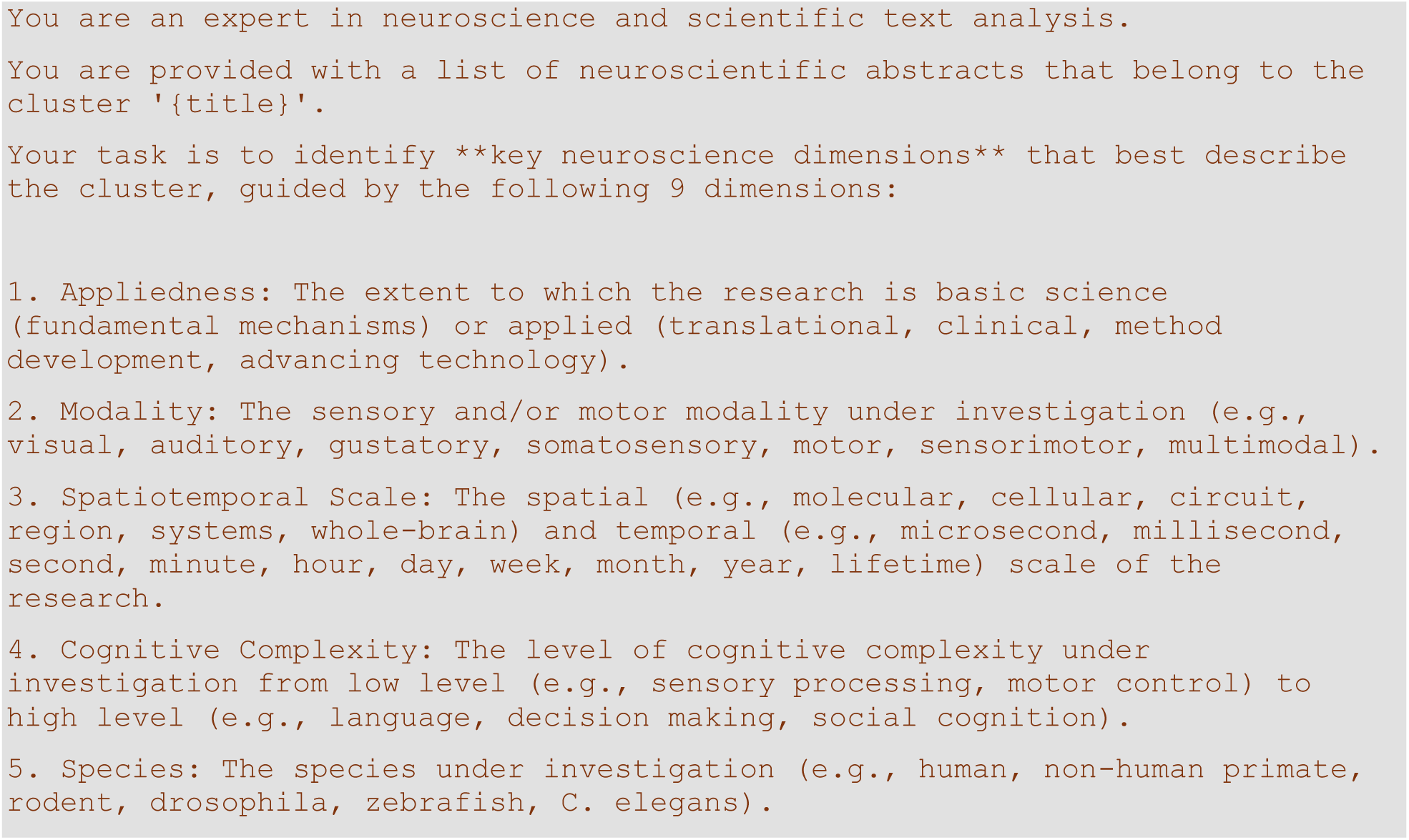
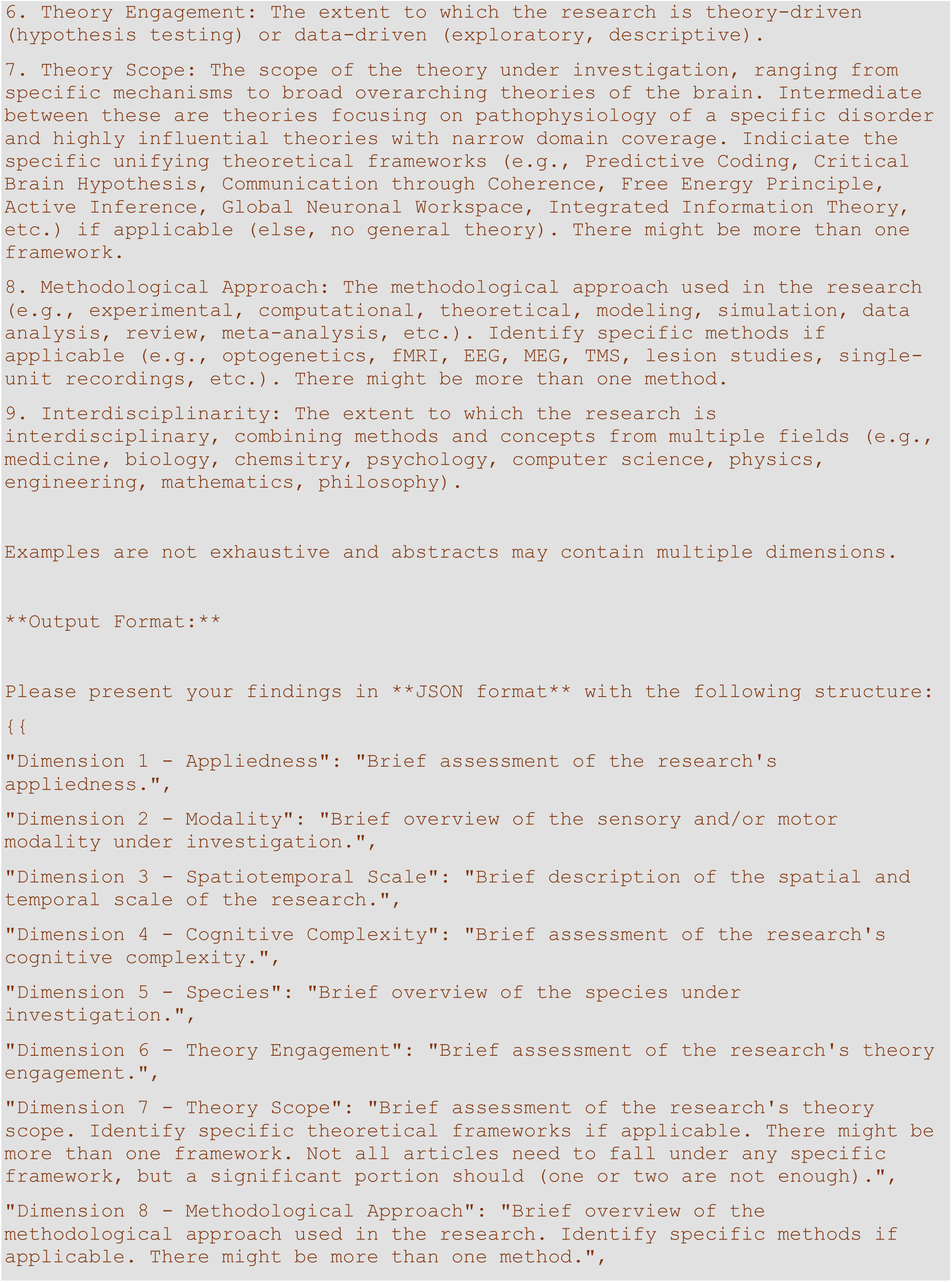
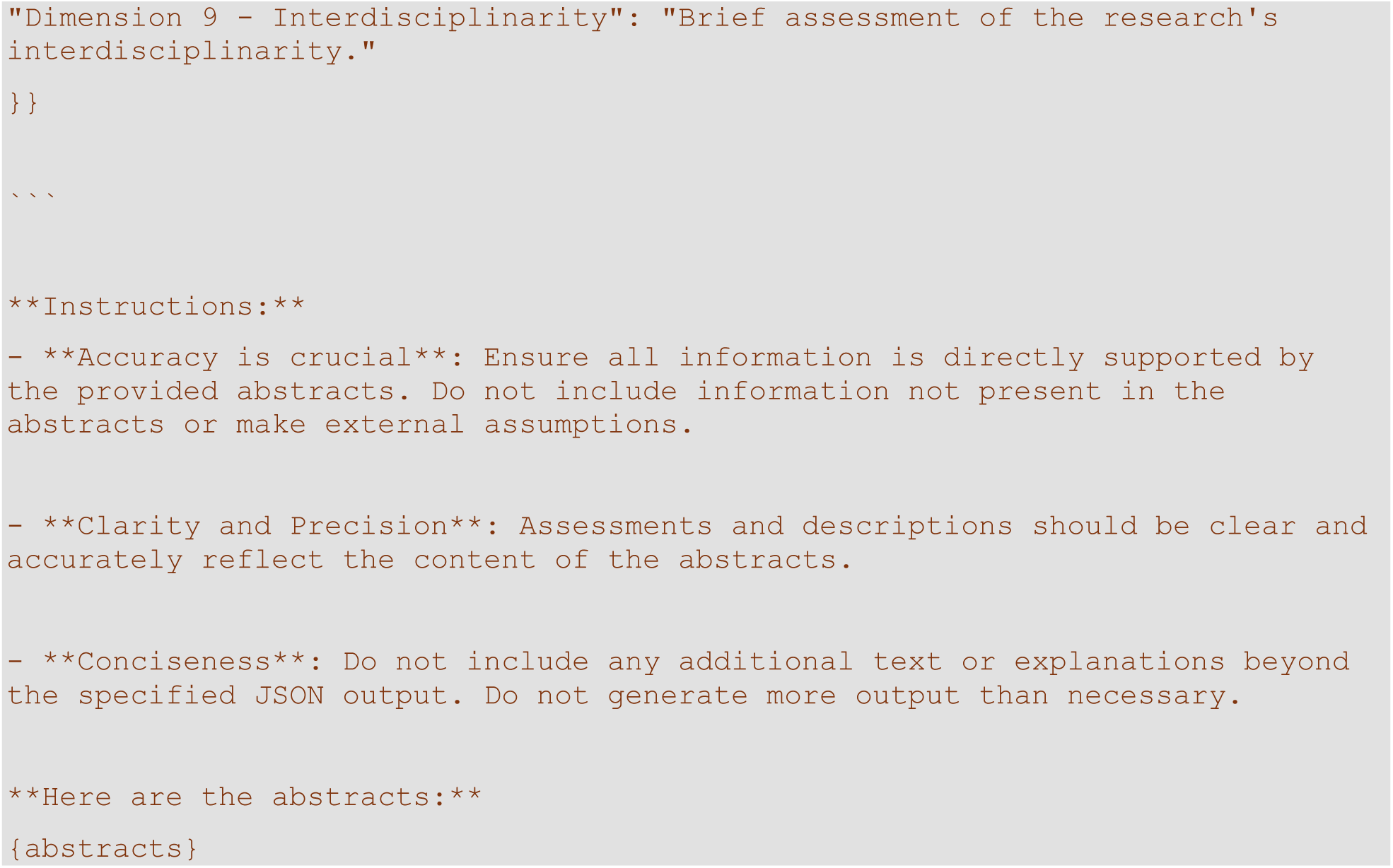

The generated JSON data was automatically extracted and integrated into the dataset. It was also used for a second-level assessment of research dimensions along specific categories within each dimension. For this analysis, I split spatiotemporal scale into two dimensions, spatial scale and temporal scale. To obtain a binary judgment whether a cluster qualifies for a particular category, I submitted the verbal assessment of a cluster’s dimensions to the LLM together with specific instructions to evaluate a particular category belonging to a particular dimension. I used the following prompt for this purpose:

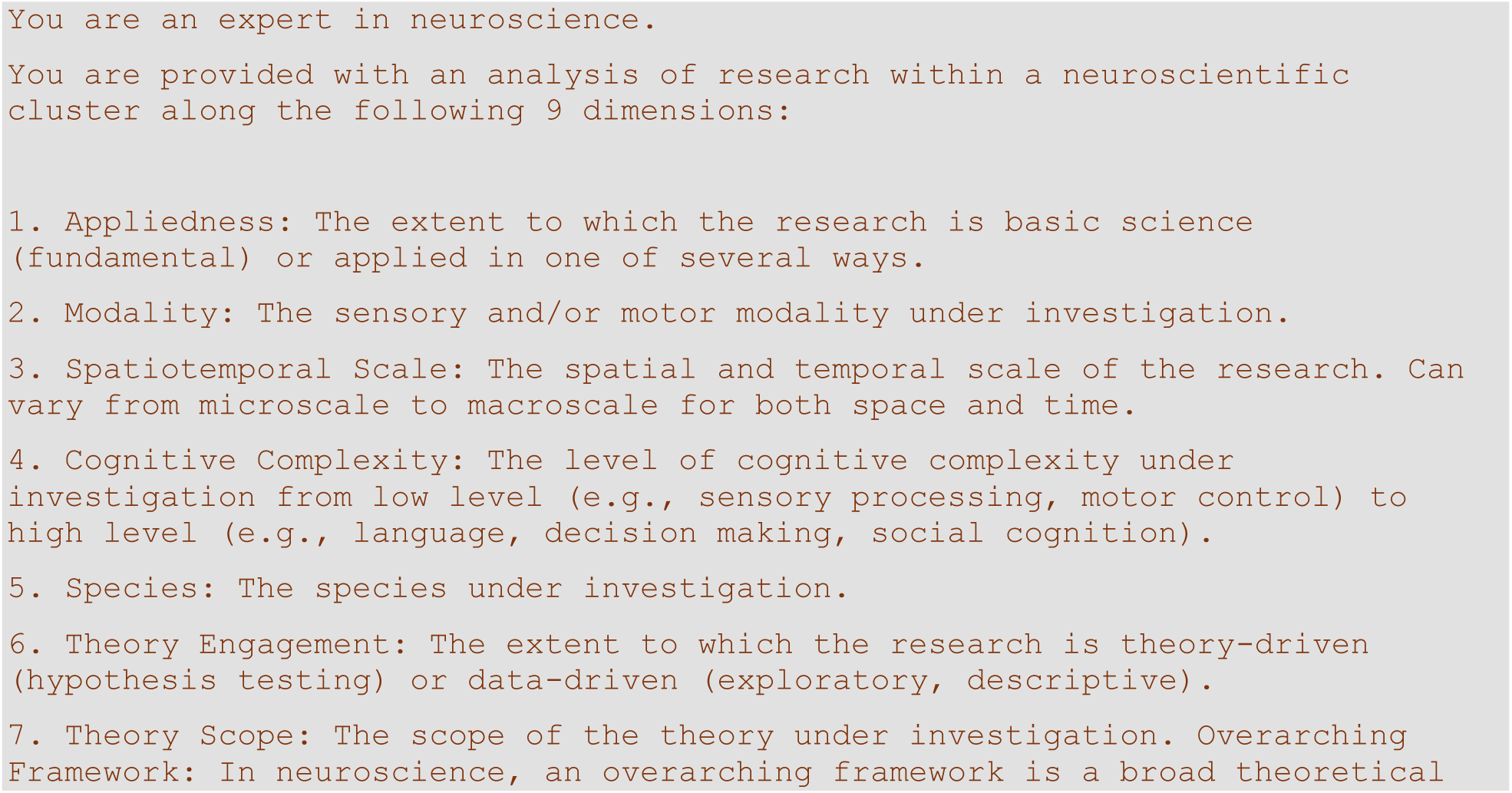
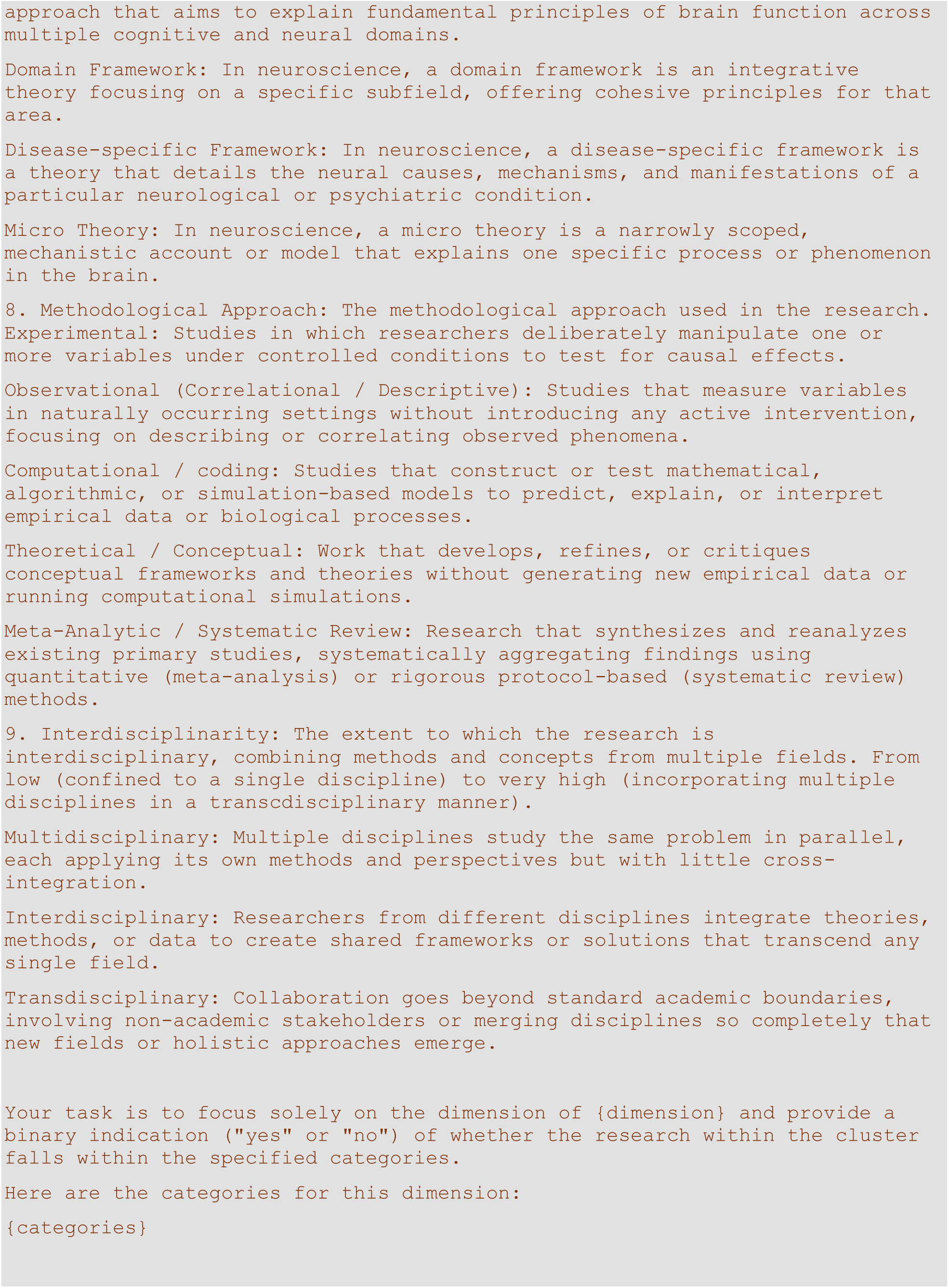
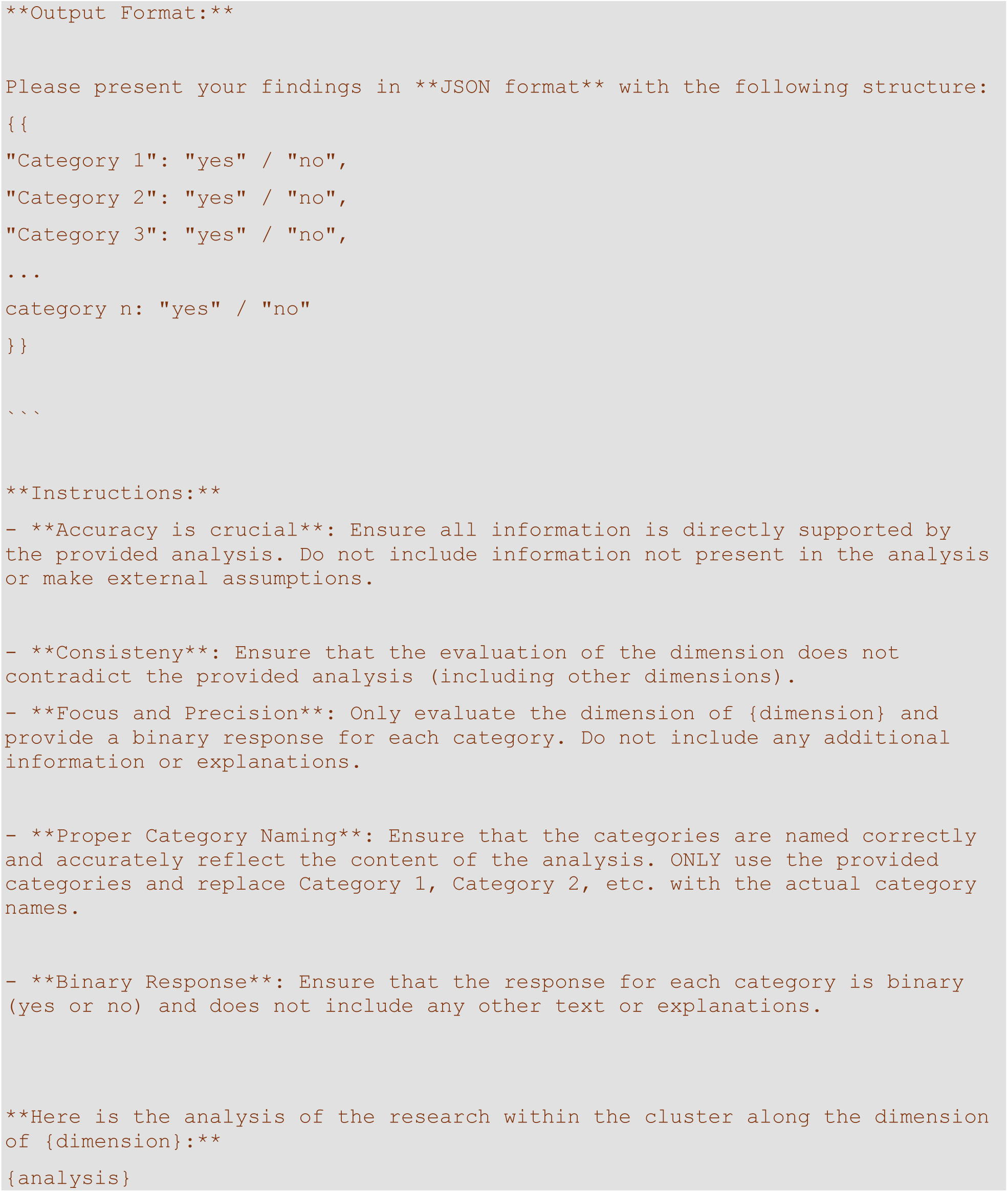

The generated JSON data was automatically extracted and integrated into the dataset.

Next, I used the LLM to analyze emerging trends in the field. To that end, I submitted abstracts of up to 200 randomly selected articles published between 2010 and 2021 labeled as older abstracts as well as abstracts of up to 200 articles published since 2021 labeled as younger abstracts to the LLM. For younger articles, I selected articles based on their citation rate. For clusters with more than 200 articles published from 2021 this means I selected the 200 most impactful articles. The LLM was instructed to identify emerging themes and methodological approaches as well as declining themes and methodological approaches using the following prompt:

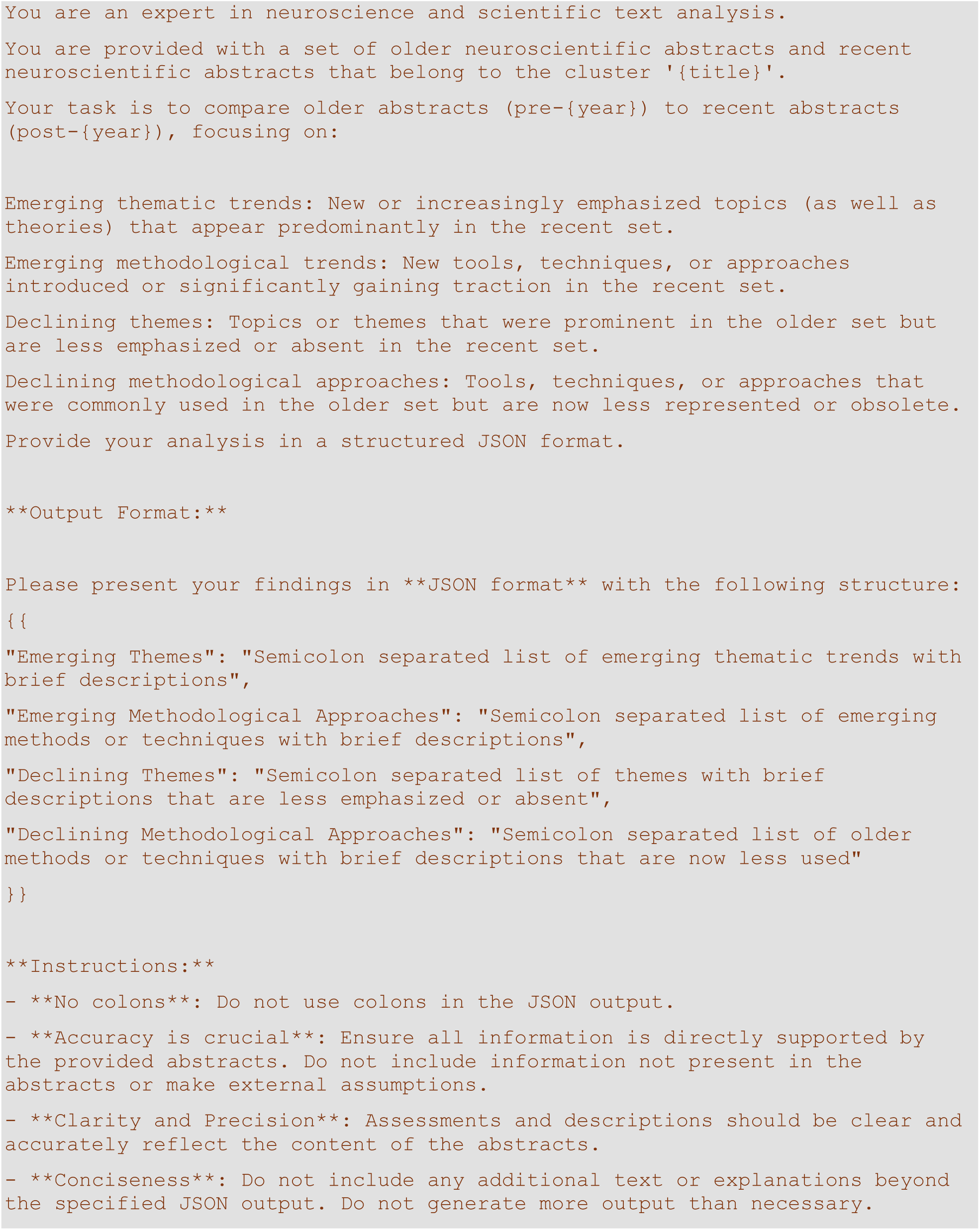
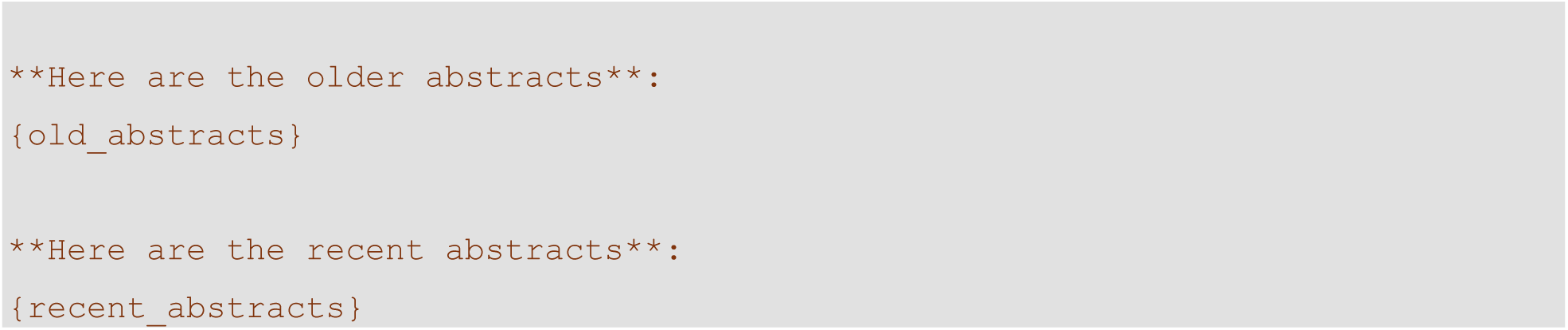

To further understand developments within each cluster, I used the LLM to extract open questions for each neuroscientific cluster. To that end I obtained five full review articles per cluster in portable document format (PDF). These review articles were published either in 2023 or 2024. Those published in 2023 were directly selected from the dataset based on their abstracts’ similarity to cluster centroid and whether I had access. Candidates for review articles published in 2024 for each cluster were identified by a PubMed query with the cluster’s keywords. Abstracts and digital object identifiers of the top 100 responses were obtained. Abstracts were then embedded and compared to the cluster centroid. Those articles with the highest similarity to which I had access were selected. I included three articles from the dataset (published in 2023) and two articles not in the dataset (published in 2024) unless access limitations required adjustments to this distribution. The list of review articles I used is shown in Supplementary Table 5. The set of five review articles of a cluster were then submitted to the LLM with the instructions to identify the major open questions that are explicitly or implicitly discussed in these articles defined in the following prompt:

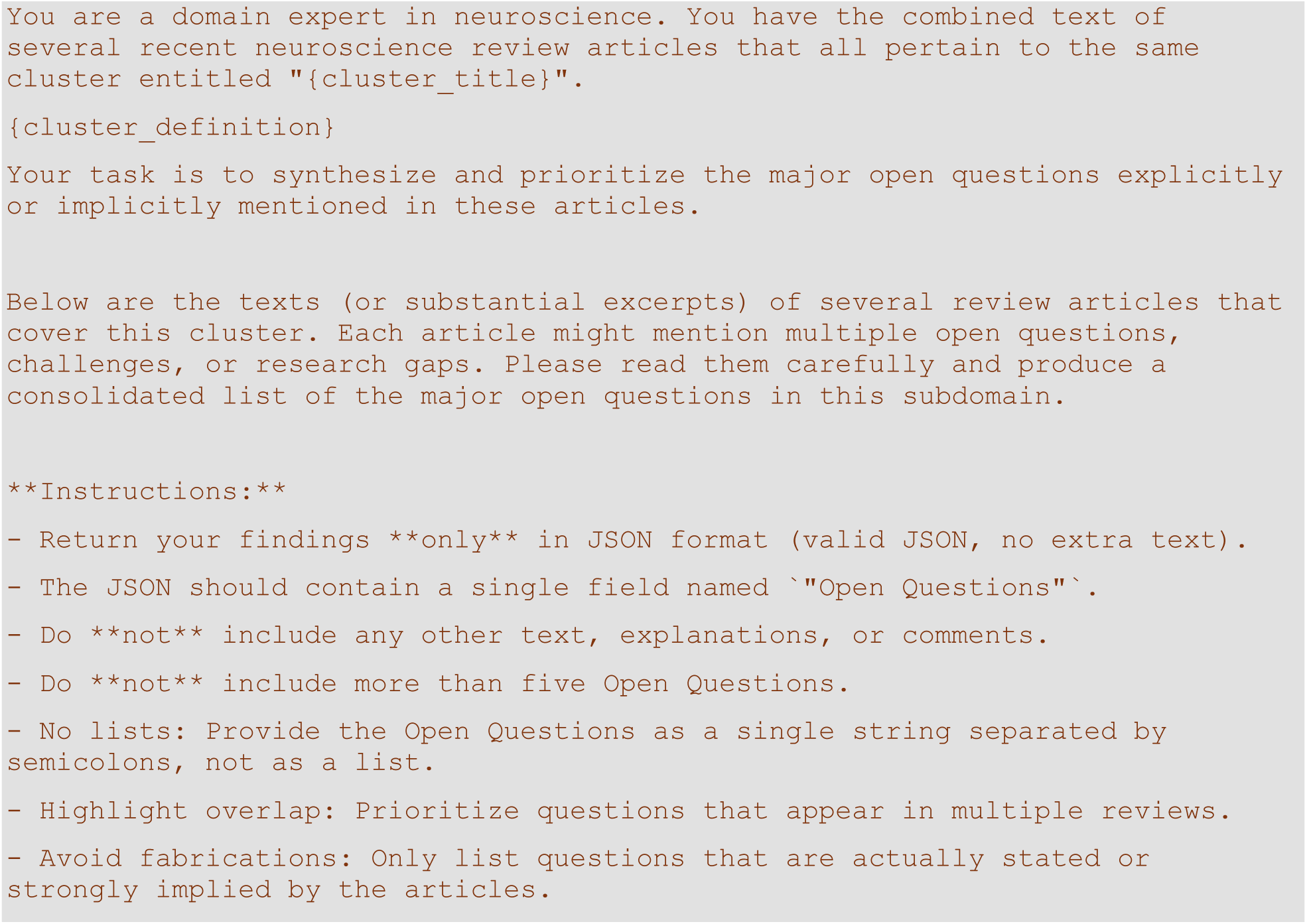
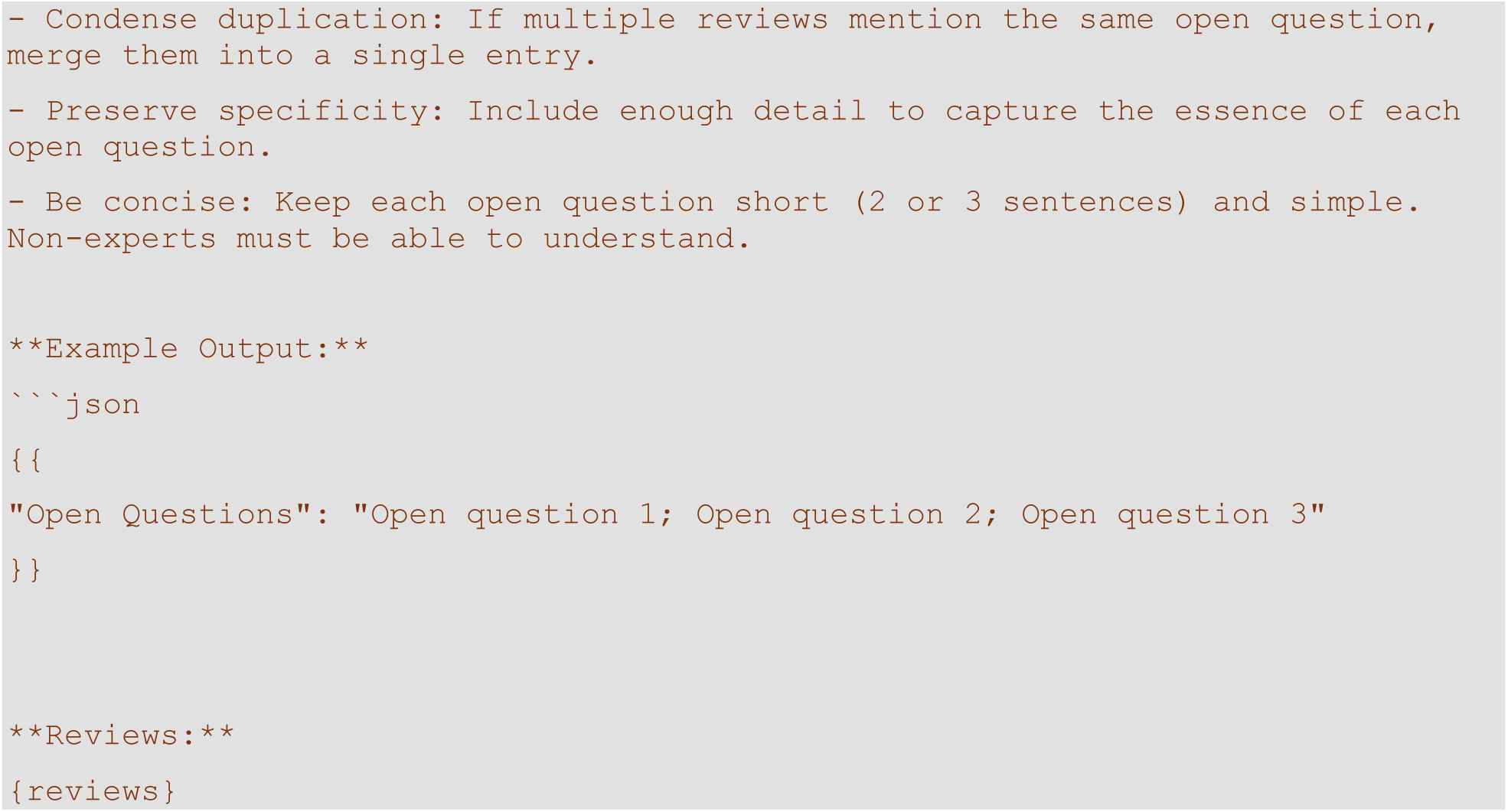

To synthesize overarching trends and key open questions that transcend individual research clusters, I systematically aggregated the results obtained from cluster-specific analyses. Each cluster’s trends and open questions were concatenated into a single document and submitted to the LLM with the instruction to identify patterns that generalize across multiple clusters, distinguishing between emerging research directions, methodological developments, and critical gaps in the field. I used the following prompt:

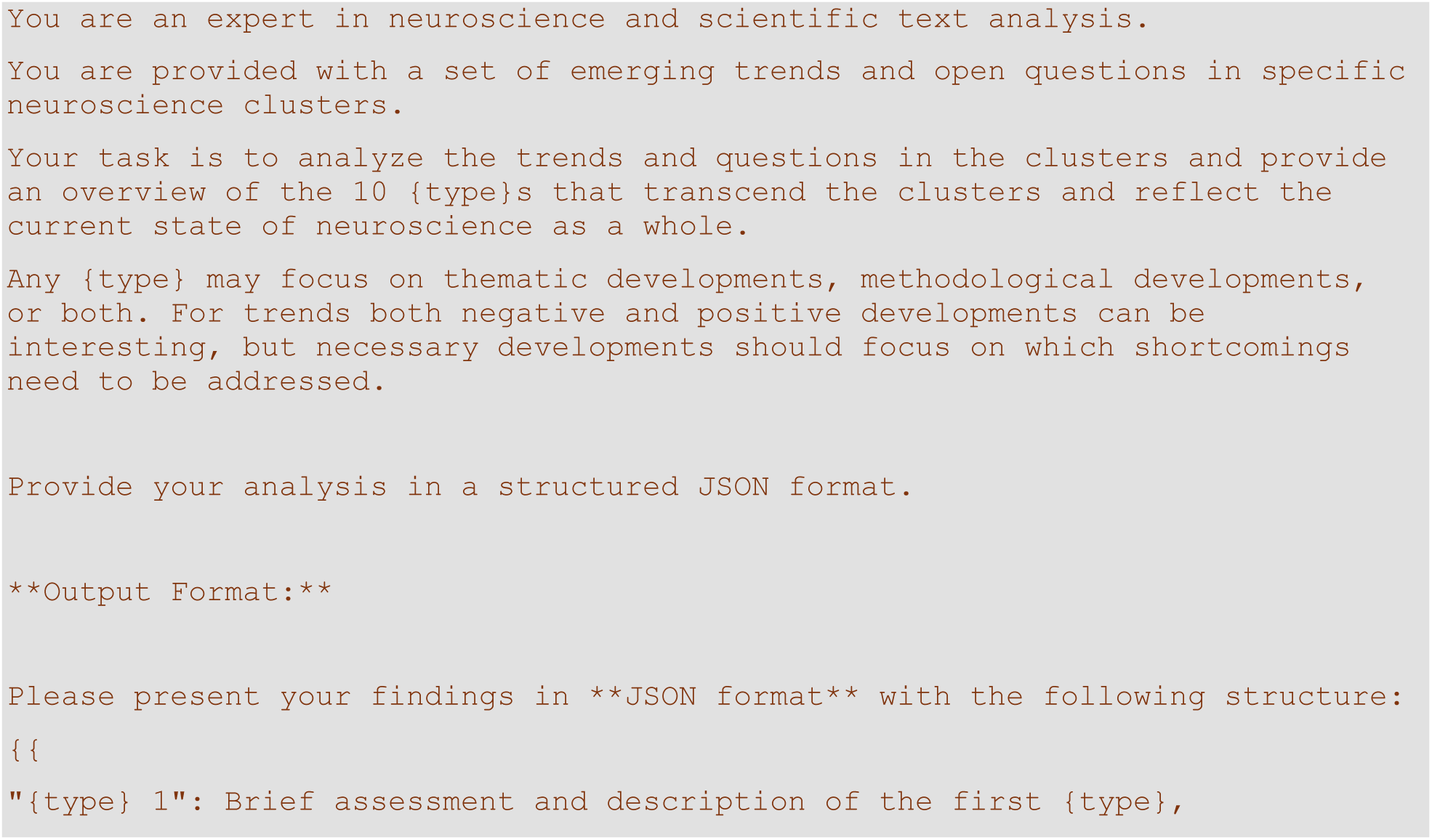
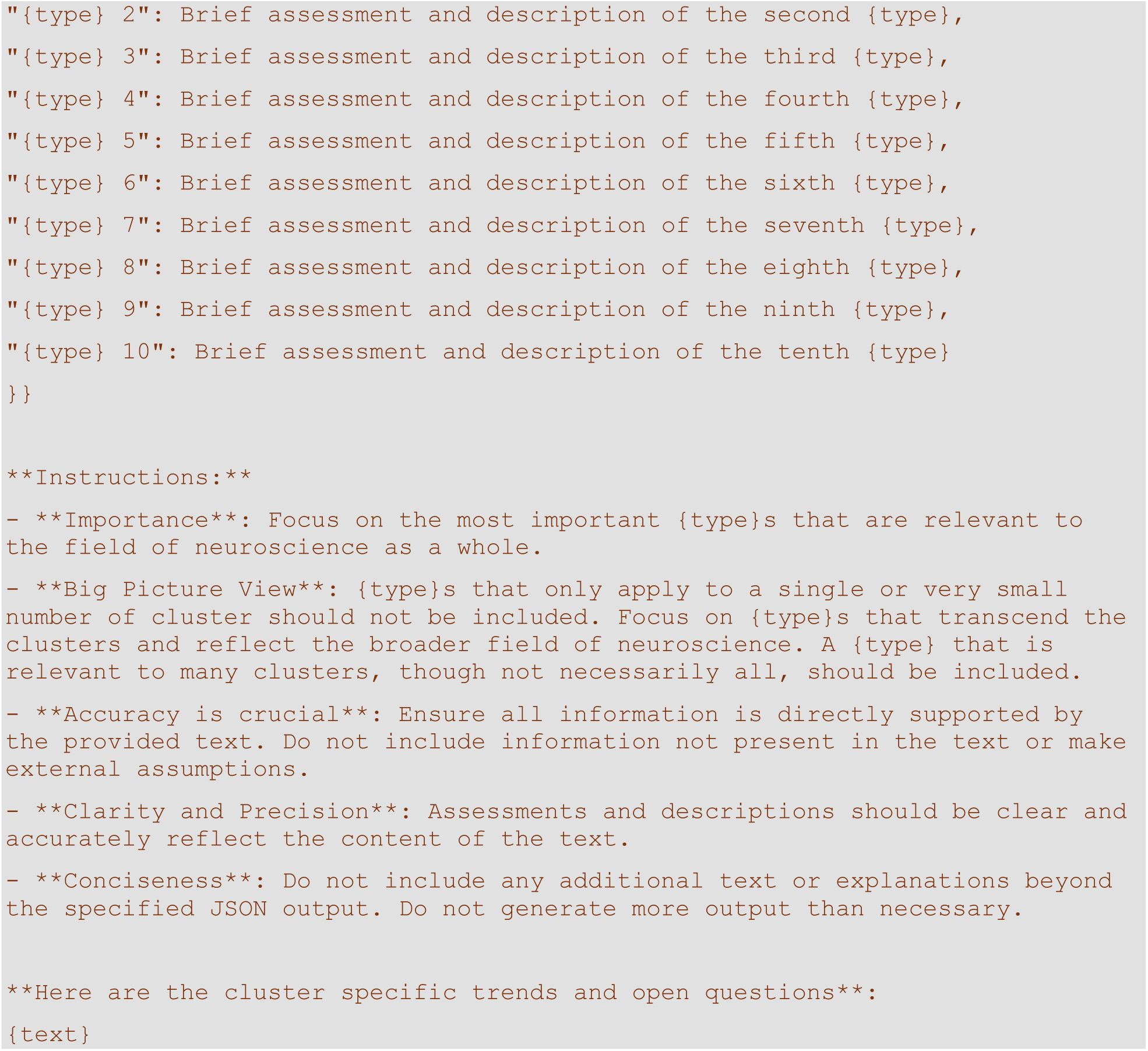

As can be appreciated from the prompt, the particular analysis the LLM should perform is variable. The prompt was constructed in this way to enable separate analyses focusing on (1) overarching trends, (2) necessary developments, and (3) transcendental research questions. In each case, the LLM was prompted to generate exactly ten items that encapsulate broad, field-wide patterns rather than localized trends specific to a single or small subset of clusters.

### LLM-Based Writing Assistance

I utilized OpenAI’s o1 (o1-2024-12-17) large language model to enhance the articles’ language and readability by submitting paragraphs together with the following prompt:

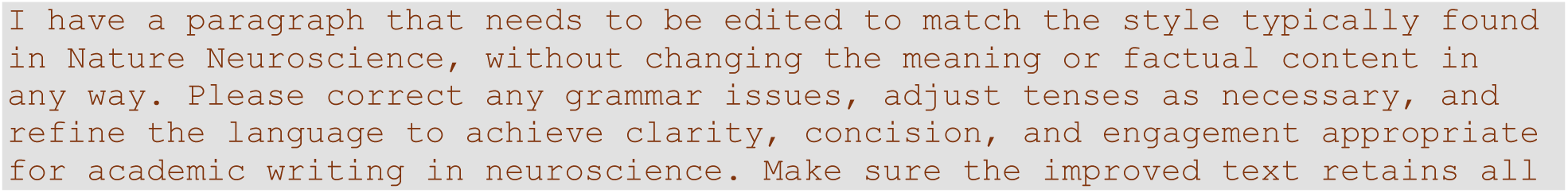
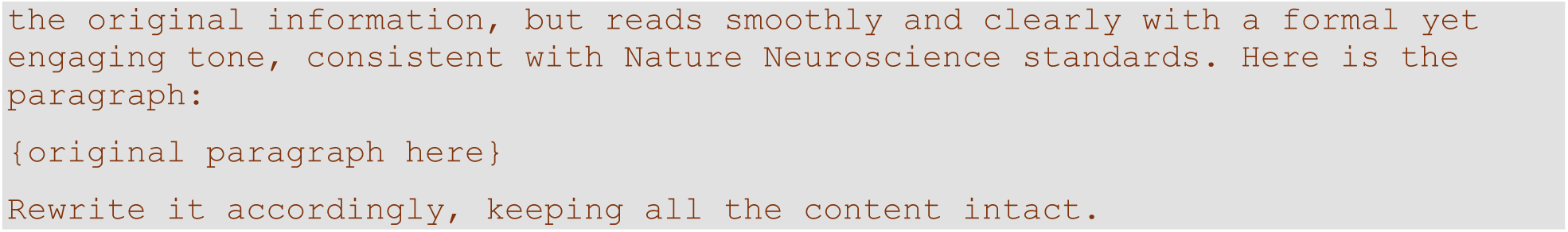

I carefully reviewed, edited, and revised the response and take ultimate responsibility for the content of this work.

## Supporting information

Supplementary Table 1: Overview of Neuroscience Domain Clusters.

Supplementary Table 2: Citation Network Metrics for Neuroscience Domain Clusters.

Supplementary Table 3: Characterization of Neuroscience Domain Clusters Across Key Dimensions

Supplementary Table 4: Emerging and Declining Trends in Neuroscience Domain Clusters

Supplementary Table 5: Bibliography of Review Articles.

## Data Availability

The data described in this study are available on Zenodo at https://doi.org/10.5281/zenodo.14865161

## Code Availability

All code is available at https://github.com/ccnmaastricht/NeuroScape.git

## Acknowledgments

I thank Arie H. van der Lugt and Gorka Zamora-Lopez for their helpful comments on earlier versions of the manuscript.

## Supplementary Materials

### Tables

***Supplementary Table 1: Overview of Neuroscience Domain Clusters*.**

*This table presents a structured summary of the 175 research clusters identified through text embedding and clustering analysis of neuroscientific literature from 1999 to 2023. Each cluster is characterized by a unique identifier, the number of articles it contains, a descriptive title, and a detailed summary of its thematic content. The table also includes a set of relevant keywords that capture the core topics and methodologies associated with each cluster*.

***Supplementary Table 2: Citation Network Metrics for Neuroscience Domain Clusters*.**

*This table presents key network metrics describing the citation structure of each research cluster, including Krackhardt coefficients for references and citations, in-degree (number of citing clusters), out-degree (number of referenced clusters), betweenness centrality (a measure of a cluster’s role as a bridge in the citation network), and PageRank (an indicator of a cluster’s relative influence based on citations from other influential clusters)*.

***Supplementary Table 3: Characterization of Neuroscience Domain Clusters Across Key Dimensions*.**

*This table provides a systematic classification of each research cluster along nine fundamental dimensions of neuroscientific inquiry, including appliedness, modality, spatiotemporal scale, cognitive complexity, species studied, theory engagement, theory scope, methodological approach, and interdisciplinarity. Note that spatiotemporal scales are divided into spatial and temporal scales in the manuscript*.

***Supplementary Table 4: Emerging and Declining Trends in Neuroscience Domain Clusters*.**

*This table summarizes the evolving thematic and methodological landscape within neuroscience research clusters. It highlights emerging research themes and methodological approaches, as well as declining areas of focus. Additionally, key open questions are listed for each cluster, reflecting unresolved scientific challenges and potential future research directions*.

***Supplementary Table 5: Bibliography of Review Articles*.**

